# Differential DNA damage response to WRN inhibition identifies a targetable vulnerability in ARID1A-mutated cancers

**DOI:** 10.1101/2025.05.04.652100

**Authors:** Jiwon Kim, Jaeik Oh, Dongjun Jang, Seungjae Shin, Soo-Jin Lee, Sang Eun Lee, Yoojin Yang, Dohee Kim, Ae-Jin Choi, Hae Rim Jung, Yumi Oh, Sung-Yup Cho

## Abstract

ARID1A, a key subunit of the SWI/SNF chromatin remodeling complex, is frequently mutated in cancers. However, effective clinical treatments for patients with this mutation are limited, highlighting a demand for new therapeutic strategies. Here, we establish Werner syndrome ATP-dependent helicase (WRN) as a critical vulnerability target in ARID1A-mutated cancers. Upon genetic and pharmacological inhibition of WRN, ARID1A-mutated cells had defective Chk1-mediated DNA damage signaling, resulting in compensatory Chk2 activation, leading to G1-phase arrest and apoptosis, whereas ARID1A-proficient cells underwent Chk1-dependent G2/M arrest. Additional p21 inhibition, combined with WRN suppression, drove G1-arrested cells to re-enter the cell cycle, triggering mitotic catastrophe. The anti-tumor efficacy of WRN inhibition was validated using in vivo cell lines and patient-derived xenograft mouse models. Our findings define WRN as a selective therapeutic target in ARID1A-mutated cancers and suggest a combinatorial strategy of WRN and p21 inhibition as a therapeutic approach.

## Introduction

Chromatin remodeling is a molecular process of modifying chromatin structure to regulate gene expression, and its dysregulation is associated with carcinogenesis. ^1^ ARID1A (AT-rich interaction domain 1A), a core subunit of the SWI/SNF chromatin remodeling complex, is frequently mutated in many cancer types; its mutation accounts for approximately 6% of all cancer cases. ^2^ ARID1A mutations are particularly prevalent in colorectal cancer, ^3^ endometrial cancer, ^4^ ovarian clear cell cancer, ^5^ and gastric cancer. ^6^ The high incidence of loss-of-function mutations in the *ARID1A* gene poses a major challenge for the development of targeted therapies in ARID1A-mutated cancers. To address this, several synthetic lethal targets have been suggested for ARID1A-mutated cancers, including ARID1B; ^7^ EZH2; ^8^ PARP; ^9^ ATR; ^10^ HDAC6, ^11^ and AURKA-CDC25C. ^12^ However, the clinical application of synthetic lethality-based therapeutics in ARID1A-mutant cancers remains limited, underscoring the need for new therapeutic strategies.

Recently, WRN has been identified as a synthetic lethal target in microsatellite instability (MSI) cancers. ^13^ WRN is a RecQ-like DNA helicase that plays a major role in the restart of homologous recombination-mediated replication forks and in preventing fork collapse. ^14^ WRN depletion selectively induces cell death in MSI cancer cells, with loss of the helicase domain rather than the exonuclease domain. ^13^ Moreover, WRN silencing leads to double-strand break (DSB) accumulation, activating DSB repair pathways and resulting in cell cycle arrest at either the G1 or G2/M phases in MSI cancer cells. ^13^ However, the precise mechanism governing the differential cell cycle arrest upon WRN depletion in MSI cancer cells remains unclear. Interestingly, ARID1A mutations are frequently associated with the MSI subgroup in gastric and colorectal cancer, ^6, 15, 16^ and ARID1A deficiency has been linked to a mismatch repair-deficient phenotype, ^17^ suggesting the potential role for ARID1A mutation in MSI cancers.

In this study, we investigated WRN as a therapeutic vulnerability target in ARID1A-mutated cancers. Upon WRN depletion, defects in Chk1 signaling in ARID1A-mutated cells promoted compensatory Chk2 activation. This differential activation of DNA damage repair pathways led to a distinct pattern of cell cycle arrest, with G1 arrest being preferentially induced in ARID1A-mutated cells and G2/M arrest induced in ARID1A wild-type cells. Furthermore, additional inhibition of p21 forced G1-arrested ARID1A-mutated cells to bypass cell cycle arrest, ultimately leading to mitotic catastrophe. Our findings suggest that WRN depletion impairs the viability of ARID1A-mutated cancer cells by altering DNA damage responses and inducing distinct cell cycle arrest. Additionally, we found that dual targeting of p21 and WRN further enhanced cell death by promoting mitotic catastrophe, providing a potential combination treatment strategy for ARID1A-mutated cancers.

## Results

### Identification of WRN as a therapeutic vulnerability in ARID1A-mutated MSI cancers

To investigate the new therapeutic targets of ARID1A-mutated cancers, we analyzed the Project Achilles dataset, which contains genetic dependency values (β-scores) for each gene across the Cancer Cell Line Encyclopedia (CCLE). ^18^ Cell lines were classified based on *ARID1A* mutation status, and β-scores for each gene were compared between ARID1A wild-type and ARID1A-mutant cell lines (Fig. 1a). Differential dependency analysis using limma identified 50 genes with significantly altered dependencies in ARID1A-mutant cell lines (|mean β-score difference| > 0.1, FDR q < 0.05) (Fig. 1b). Gene set analysis using the ClueGO plugin for REACTOME Pathway gene sets revealed that differentially dependent genes were associated with “DNA damage response”, “cell cycle progression (G1 and G2/M)”, and “tricarboxylic acid cycle” (Fig. 1c). Additionally, ClueGO analysis with Gene Ontology Biological Process gene sets identified enrichment in pathways such as “regulation of G0-to-G2 transition” and “centrosome cycle” (Extended Data Fig. 1a). These findings suggest that *ARID1A* mutations confer differential dependencies on genes involved in DNA damage response and cell cycle regulation.

**Fig. 1.**
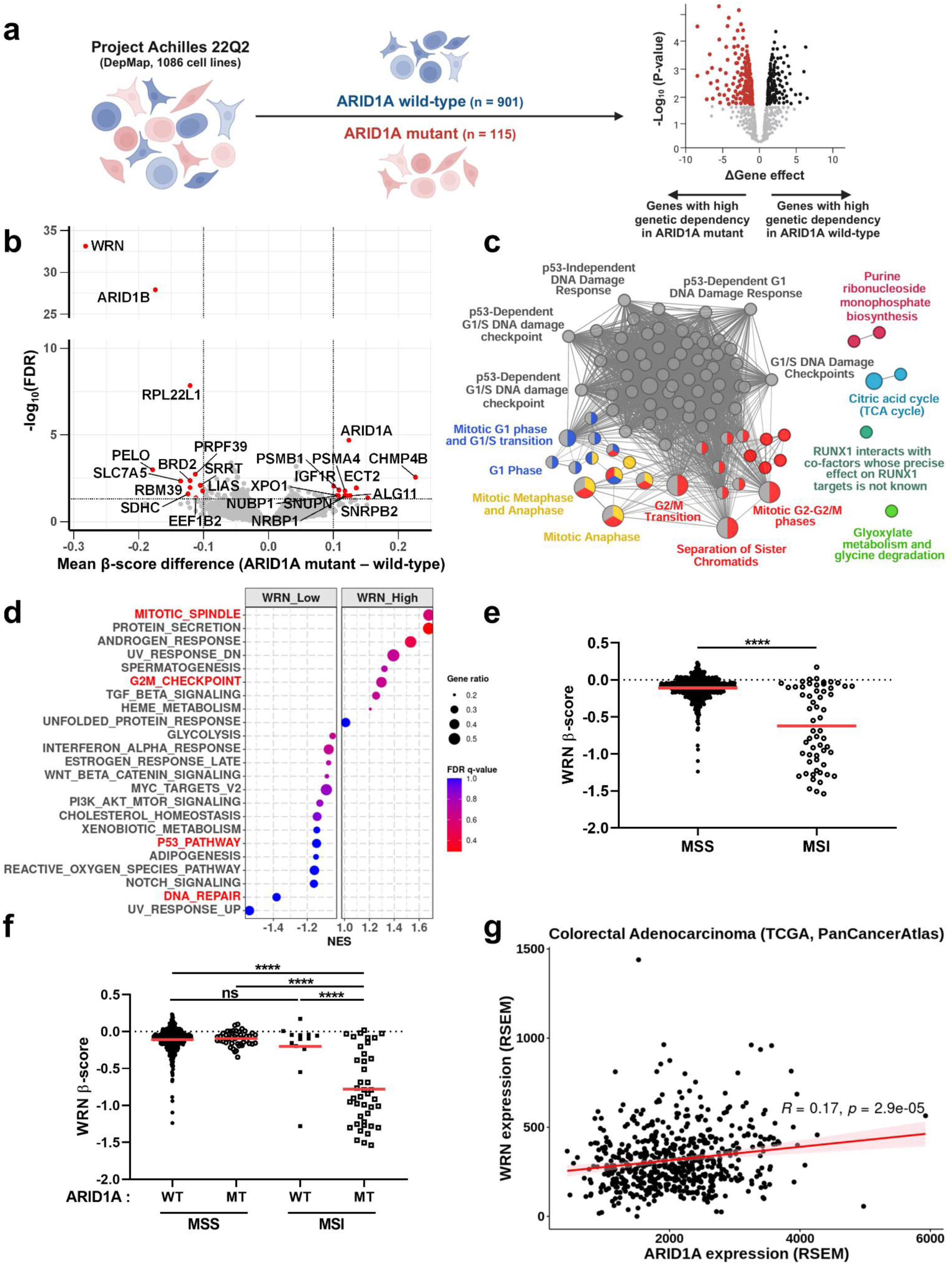
ARID1A mutation confers selective dependency on WRN in cancer cell lines. (a) Schematic representation of the analysis using Project Achilles 22Q2 data from DepMap, stratified by ARID1A-mutation status. (b) Volcano plot showing false discovery rate (FDR) q-values against the difference of mean dependency scores between ARID1A wild-type (n = 901) and ARID1A-mutant (n = 115) cell lines from Project Achilles. Red dots represent genes with |mean β-score difference| > 0.1 and FDR q < 0.05, while gray dots represent all other genes. FDR q-values were calculated using the Benjamini–Hochberg method. (c) ClueGO enrichment analysis of differentially essential genes based on REACTOME Pathway gene sets. Nodes represent REACTOME terms, with node size indicating enrichment significance. Functionally related terms are connected by edges, and distinct functional categories are color-coded and labeled. (d) Gene set enrichment analysis comparing WRN-high and WRN-low expression groups in The Cancer Genome Atlas colorectal adenocarcinoma datasets (n = 296 for each group) using hallmark gene sets. Gene sets are displayed with normalized enrichment scores (NES) on the x-axis. Dot size represents the proportion of core enrichment genes relative to the total genes in each gene set. Dot colors denote FDR q-values. Gene sets were only presented if |NES| > 1. (e) Comparative analysis of β-scores for WRN in microsatellite stable (MSS, n = 658) and microsatellite instable-high (MSI-H, n = 58) cell lines. A red line indicates the mean β-score for each group. Statistical significance was determined using Welch’s unpaired t-test. (f) Comparative analysis of β-scores for WRN in MSS/ARID1A wild-type (n = 575), MSS/ARID1A-mutant (n = 44), MSI-H/ARID1A wild-type (n = 13), and MSI-H/ARID1A-mutant (n = 41) cell lines. A red line indicates the mean β-score for each group. Statistical significance was determined using Tukey’s post hoc test following one-way analysis of variance (ANOVA). (g) Correlation plot of *ARID1A* and *WRN* mRNA expression levels in TCGA colorectal carcinoma datasets (n = 592). A red line represents the linear regression fit, and the light-red shaded region indicates the confidence interval. Correlation analysis was assessed using Pearson correlation analysis. RSEM, RNA-Seq by expectation maximization. Statistical significance was defined as follows: not significant; ns, p > 0.05; *, p ≤ 0.05; **, p ≤ 0.01; ***, p ≤ 0.001; ****, p ≤ 0.0001.

Among genes with differential dependencies, *WRN* had the highest genetic dependency in ARID1A-mutant cell lines, followed by *ARID1B*, which has been previously identified as a synthetic lethal target in ARID1A-mutated cancers (Fig. 1b and Extended Data Fig. 1b). ^7^ Since WRN is a member of the RecQ helicase family, we compared the β-scores of other RecQ family members; however, only *WRN* displayed a comparable increase in dependency in ARID1A-mutant cell lines (Extended Data Fig. 1b). When we performed Gene Set Enrichment Analysis (GSEA) using hallmark gene sets to compare patients with colorectal adenocarcinoma with high or low WRN expression, classified based on median WRN expression level (RSEM = 287.035) in The Cancer Genome Atlas (TCGA), the WRN-high group was enriched in the “mitotic spindle” and “G2/M checkpoint” gene sets, whereas the WRN-low group was enriched in the “p53 pathway” and “DNA repair” gene sets (Fig. 1d). Taken together, the increased *WRN* dependency in ARID1A-mutated cells suggests that WRN may serve as a new therapeutic target by modulating DNA damage response and cell cycle regulation.

WRN has also been proposed as a synthetic lethal target in MSI cancers. ^13^ To investigate the relationship between MSI status and *ARID1A* mutation in *WRN* dependency, we first stratified the Project Achilles dataset by MSI status (Extended Data Fig. 1c). Consistent with previous findings, ^13^ *WRN* β-scores were generally lower in MSI cell lines, indicating higher dependency (Fig. 1e). However, some MSI cell lines had β-scores similar to those of microsatellite stable (MSS) cell lines, highlighting variability in *WRN* dependency even in MSI cells (Fig. 1e). To identify genetic mutations, including *ARID1A* mutations, associated with *WRN* dependency, we categorized the MSI cell lines into high- and low-dependency groups using a β-score threshold corresponding to the median (β-score = −0.617) (Extended Data Fig. 1c). Mutation profiling of frequently mutated oncogenes and tumor suppressors revealed distinct mutation patterns in the *ARID1A*, *PIK3CA*, *TP53*, and *BRAF* genes between the *WRN* high- and low-dependency groups (Extended Data Fig. 1d). For genes with a distinct mutation pattern, we analyzed *WRN* β-scores based on the mutation status of each gene. Notably, MSI cell lines lacking ARID1A mutations exhibited *WRN* dependency similar to MSS cell lines, whereas high *WRN* dependency (indicated by low β-scores) was observed exclusively in ARID1A-mutated MSI cell lines (Fig. 1f). However, this trend was not observed when comparing MSI cell lines with or without mutations in *PIK3CA*, *TP53*, or *BRAF* (Extended Data Fig. 1e). These results underscore the significance of ARID1A mutations in driving *WRN* dependency in MSI cancers, supporting WRN as a potential therapeutic target, particularly in MSI and ARID1A-mutated contexts.

### ARID1A regulates WRN expression at the transcriptional level

To investigate the relationship between ARID1A and WRN, we analyzed mRNA expression data from The Cancer Genome Atlas (TCGA) and Cancer Cell Line Encyclopedia (CCLE). Correlation analysis revealed a positive correlation between *ARID1A* and *WRN* expression (colorectal adenocarcinoma, TCGA, R = 0.17; all cell lines in CCLE, R = 0.4; large and small intestine cell lines in CCLE, R = 0.22) (Fig. 1g and Extended Data Fig. 2a). In isogenic HCT116 human colon cancer cell models with ARID1A mutations (ARID1A wild-type, ARID1A-Q456*/+, ARID1A-Q456*/Q456*), ARID1A loss due to genetic mutation was associated with a progressive reduction in WRN expression at both the RNA and protein levels in ARID1A-mutated cells (Extended Data Fig. 2b). Similarly, analysis of public RNA sequencing data from the Gene Expression Omnibus (https://www.ncbi.nlm.nih.gov/geo, GSE101966) comparing ARID1A wild-type and ARID1A-knockout HCT116 cells confirmed *WRN* downregulation in ARID1A-knockout cells (Extended Data Fig. 2c). To assess whether ARID1A transcriptionally regulates *WRN* expression, we performed a nascent RNA synthesis assay using EU-labeled RNA, which revealed reduced transcription of *WRN* in ARID1A-mutated cells (Extended Data Fig. 2d). Consistently, *ARID1A* knockdown using siRNA in ARID1A wild-type HCT116 cells led to decreased WRN mRNA and protein levels (Extended Data Fig. 2e), while ARID1A overexpression in ARID1A-mutated cells restored WRN expression (Extended Data Fig. 2f). As ARID1A is a core subunit of the chromatin remodeling complex SWI/SNF, ^19, 20^ we hypothesized that ARID1A directly regulates *WRN* expression at the transcriptional level. ChIP-qPCR analysis confirmed that ARID1A binds directly to *WRN* promoter regions, and ARID1A loss was associated with diminished recruitment of phosphorylated RNA polymerase II (Ser5), a marker for active transcription initiation, at these sites (Extended Data Fig. 2g). Taken together, these findings suggest that ARID1A regulates *WRN* transcription by binding to its promoter region.

### WRN depletion significantly inhibits cancer cell growth in ARID1A-mutated cancers

To validate *WRN* dependency in ARID1A-mutated cancers, we used ARID1A-mutated isogenic HCT116 cell lines (ARID1A wild-type, ARID1A-Q456*/+, ARID1A-Q456*/Q456*). Consistent with previous findings, phosphorylation of Akt and S6K was enhanced in ARID1A-mutated cells (Extended Data Fig. 2h). ^21^ siRNA-mediated WRN depletion in ARID1A-mutated cells resulted in a more pronounced suppression of proliferation compared with wild-type cells, leading to the slowest growth rate (Fig. 2a and Extended Data Fig. 3a). BrdU incorporation and colony formation were also significantly reduced in ARID1A-mutated cells upon *WRN* knockdown (Fig. 2b,c). Furthermore, WRN depletion induced the highest levels of apoptosis in ARID1A-mutated cells (Fig. 2d). To further validate these effects, we generated ARID1A-knockout cells using the CRISPR/Cas9 system. Consistently, WRN knockdown in ARID1A-knockout cells resulted in a significant decrease in cell proliferation, BrdU incorporation, and colony formation, along with an increase in apoptosis, compared to HCT116 control cells (Extended Data Fig. 3b-f). Notably, ARID1A-knockout cells with sgARID1A-1 failed to form colonies. In contrast, restoration of ARID1A expression in ARID1A-mutated cells rescued cell proliferation, BrdU incorporation, and colony formation while reducing apoptosis following *WRN* knockdown, ultimately enhancing overall cell viability (Fig. 2e-h and Extended Data Fig. 3g). These findings demonstrate that WRN is essential for the viability and proliferation of ARID1A-mutated cancer cells.

**Fig. 2.**
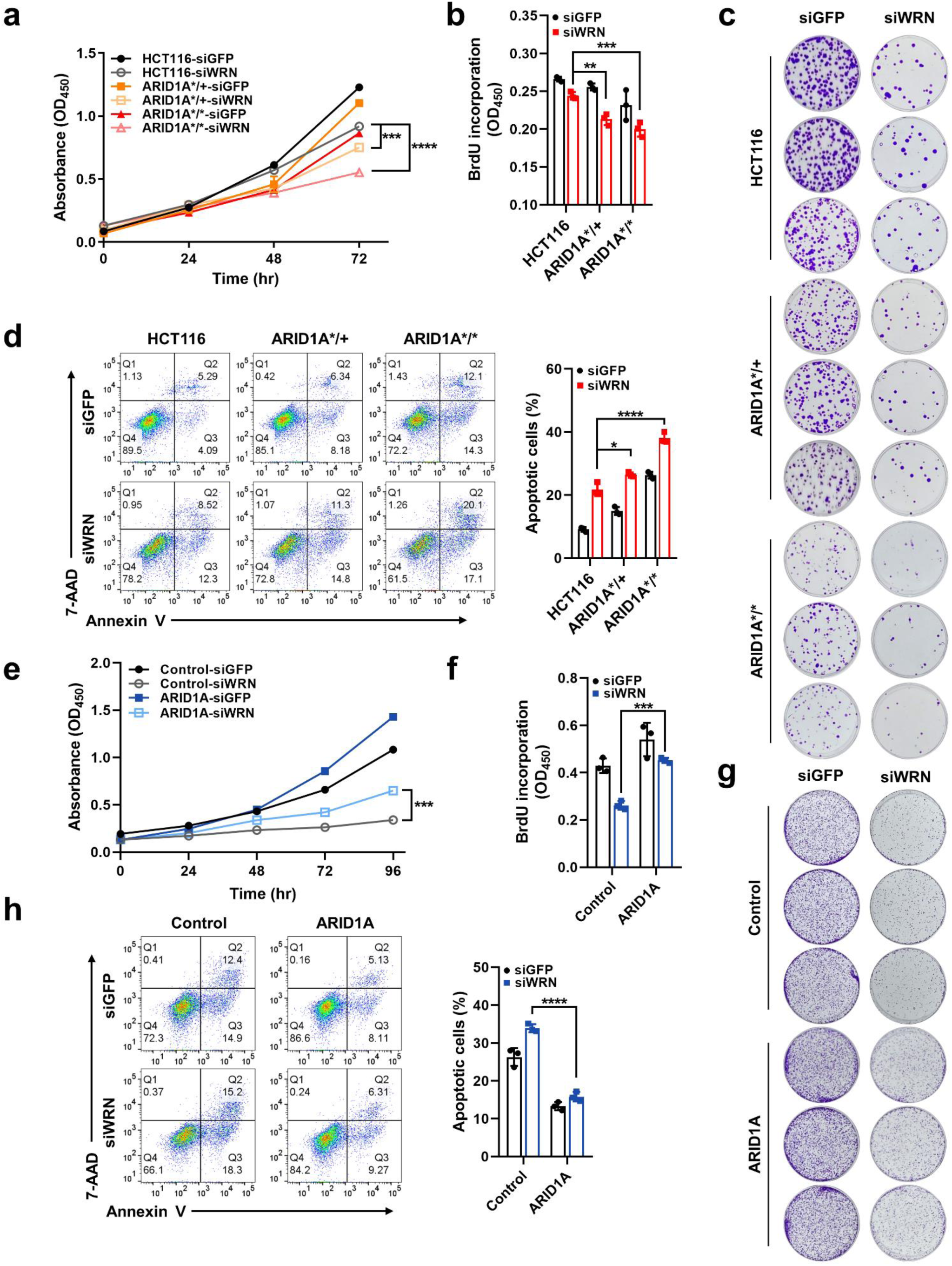
WRN depletion impairs proliferation and induces apoptosis in ARID1A-mutated cancer cells. (a) Cell proliferation measured using the WST-1 assay in ARID1A wild-type and ARID1A-mutated cells following *WRN* knockdown, measured at 0, 24, 48, and 72 hours after transfection. Data represent three independent experiments and are presented as mean ± standard deviation (SD). Statistical significance was determined using Tukey’s post hoc test following one-way analysis of variance (ANOVA). (b) BrdU incorporation assay to assess DNA synthesis in ARID1A wild-type and ARID1A-mutated cells following *WRN* knockdown. Cells were incubated with BrdU for 8 hours at 24 hours after transfection. Data represent three independent experiments and are presented as mean ± SD. Statistical significance was determined using Tukey’s post hoc test following one-way ANOVA. (c) Representative images of the colony formation assay in ARID1A wild-type and ARID1A-mutated cells following *WRN* knockdown, assessed after 9 days of culture. (d) Apoptosis assay in ARID1A wild-type and ARID1A-mutated cells following *WRN* knockdown, measured by annexin V/7-AAD staining at 72 hours after transfection. Representative flow cytometry plots (left) and quantification (right) are shown. Data represent three independent experiments and are presented as mean ± SD. Statistical significance was determined using Tukey’s post hoc test following one-way ANOVA (e) Cell proliferation measured using the WST-1 assay in control and ARID1A-overexpressing ARID1A-mutated cells following *WRN* knockdown, measured at 0, 24, 48, 72, and 96 hours after transfection. Data represent three independent experiments and are presented as mean ± SD. Statistical significance was determined using Tukey’s post hoc test following one-way ANOVA. (f) BrdU-incorporation assay to assess DNA synthesis in control and ARID1A-overexpressing ARID1A-mutated cells following *WRN* knockdown. Cells were incubated with BrdU for 8 hours at 24 hours after transfection. Data represent three independent experiments and are presented as mean ± SD. Statistical significance was determined using Tukey’s post hoc test following one-way ANOVA. (g) Representative images of the colony formation assay in control and ARID1A-overexpressing ARID1A-mutated cells following *WRN* knockdown, assessed after 9 days of culture. (h) Apoptosis assay in control and ARID1A-overexpressing ARID1A-mutated cells following *WRN* knockdown, measured by annexin V/7-AAD staining at 72 hours after transfection. Representative flow cytometry plots (left) and quantification (right) are shown. Data represent three independent experiments and are presented as mean ± SD. Statistical significance was determined using Tukey’s post hoc test following one-way ANOVA Statistical significance was defined as follows: not significant; ns, p > 0.05; *, p ≤ 0.05; **, p ≤ 0.01; ***, p ≤ 0.001; ****, p ≤ 0.0001.

### Loss of WRN activates Chk2 and induces subsequent G1 arrest in ARID1A-mutated cancer cells

To understand the molecular mechanisms underlying the pronounced anti-tumor effect of WRN depletion in ARID1A-mutated cells, we examined DNA damage signaling. WRN functions as a DNA helicase involved in critical processes such as DNA repair, replication, and telomere maintenance, ^22^ and its depletion triggers a DNA damage response in MSI cells. ^13^ In ARID1A wild-type cells, *WRN* knockdown increased Chk1 phosphorylation, whereas ARID1A-mutated cells had a relatively smaller increase in Chk1 phosphorylation but a marked elevation in Chk2 phosphorylation (Fig. 3a). Additionally, markers of DNA damage, including phosphorylated p53 and γH2AX, were also significantly elevated in ARID1A-mutated cells (Fig. 3a). Consistently, ARID1A-knockout cells had dominant Chk2 phosphorylation rather than Chk1 phosphorylation (Extended Data Fig. 4a). Reintroducing ARID1A in ARID1A-mutated cells restored Chk1 phosphorylation and reduced Chk2 phosphorylation following WRN depletion (Fig. 3b). These results suggest that ARID1A deficiency alters the DNA damage response pathway, shifting from a Chk1-dominant response to a Chk2-dominant response upon WRN depletion.

**Fig. 3.**
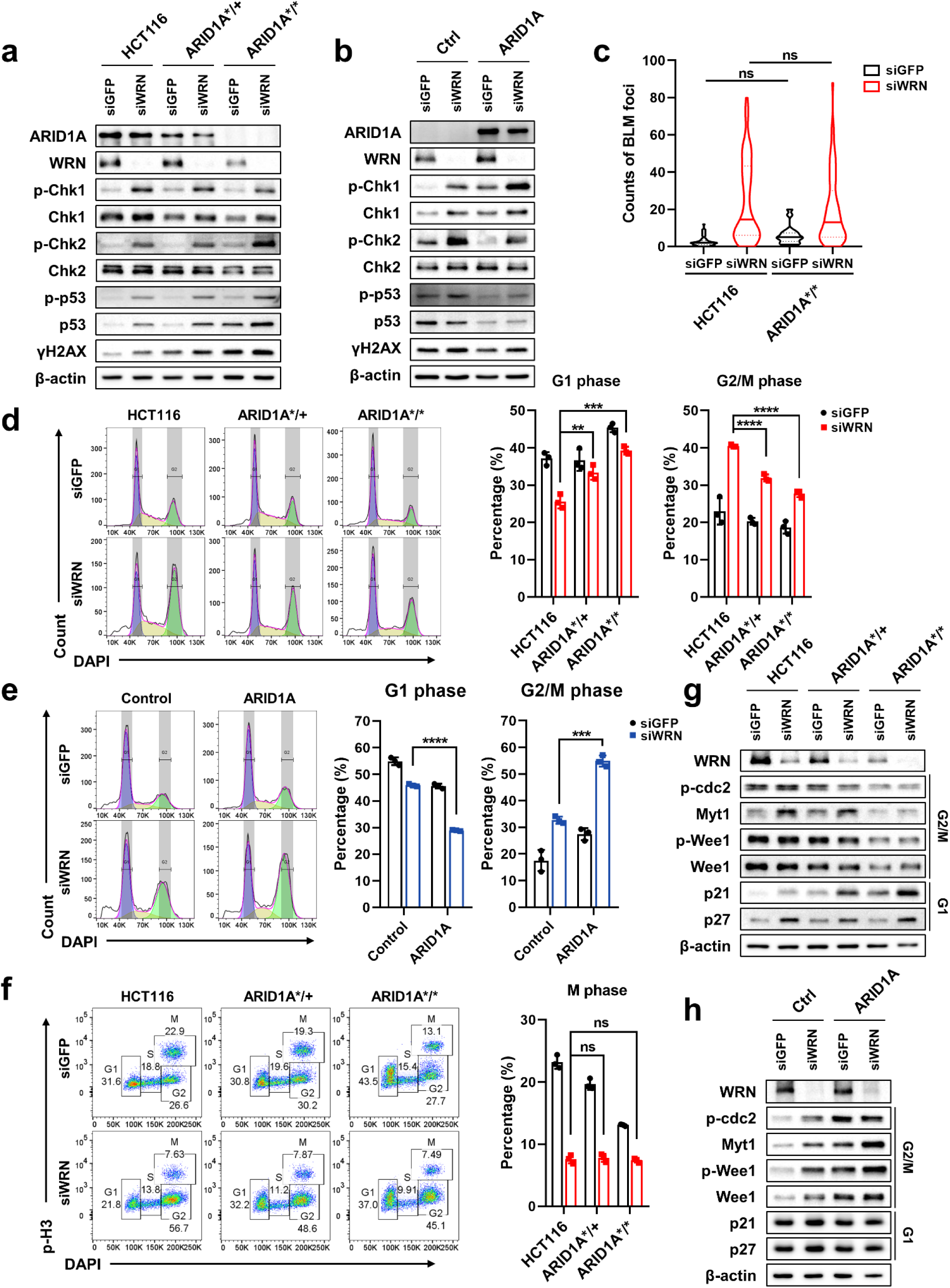
WRN depletion differentially induces DNA damage response and cell cycle alteration based on ARID1A mutation status. (a,b) Western blot analysis of DNA damage response proteins following *WRN* knockdown in ARID1A wild-type and ARID1A-mutated cells (a) and in control and ARID1A-overexpressing ARID1A-mutated cells (b). (c) Violin plot of BLM foci quantification following *WRN* knockdown in ARID1A wild-type and ARID1A-mutated cells. Foci were quantified from multiple nuclei (n = 54, 68, 34, and 71, respectively) across independent images. A solid red line represents the median, and dotted red lines indicate the quartiles. Statistical significance was determined using Welch’s unpaired t-test. (d,e) G1 and G2/M phase proportions from cell cycle analysis following *WRN* knockdown in ARID1A wild-type and ARID1A-mutated cells (d) and in control and ARID1A-overexpressing ARID1A-mutated cells (e). Representative flow cytometry plots (left) and quantification (right) are shown. Data represent three independent experiments and are presented as mean ± standard deviation (SD). Statistical significance was determined using Tukey’s post hoc test following one-way analysis of variance (ANOVA) (d) and Welch’s unpaired t-test (e). (f) Flow cytometry analysis of phospho-histone H3 (p-H3) and DAPI staining of the M phase proportion following *WRN* knockdown in ARID1A wild-type and ARID1A-mutated cells. Cells were treated with paclitaxel (10 nM, 5 hours) at 48 hours after transfection. Representative flow cytometry plots (left) and quantification (right) are shown. Data represent three independent experiments and are presented as mean ± SD. Statistical significance was determined using Tukey’s post hoc test following one-way ANOVA. (g,h) Western blot analysis of cell cycle regulatory proteins following *WRN* knockdown in ARID1A wild-type and ARID1A-mutated cells (g) and in control and ARID1A-overexpressing ARID1A-mutated cells (h). Statistical significance was defined as follows: not significant; ns, p > 0.05; *, p ≤ 0.05; **, p ≤ 0.01; ***, p ≤ 0.001; ****, p ≤ 0.0001.

Double-strand breaks primarily activate the ATM-Chk2 pathway, whereas single-strand breaks activate the ATR-Chk1 pathway. ^23, 24^ To determine whether the differential activation of Chk1 and Chk2 upon WRN depletion was linked to specific types or amounts of DNA damage, we performed comet assays. Both neutral and alkaline comet assays had comparable comet tail intensities, indicating similar types and amounts of DNA strand breaks in ARID1A wild-type and ARID1A-mutated cells following *WRN* knockdown (Extended Data Fig. 4b). Similarly, BLM helicase foci, markers for stalled replication forks, were induced at comparable levels in both cell types upon WRN depletion (Fig. 3c and Extended Data Fig. 4c), indicating that WRN depletion generates similar types and levels of DNA damage regardless of ARID1A mutation. Thus, ARID1A mutations likely influence the DNA damage response signaling pathway rather than alter the type or extent of DNA damage itself.

Chk2 arrests cells in the G1 phase via p53 stabilization in response to DNA damage, ^25^ while Chk1 mediates G2/M arrest through CDK1 phosphorylation. ^26^ To investigate whether ARID1A mutation-driven differences in Chk1 and Chk2 activation impact cell cycle regulation, we analyzed cell cycle distribution following WRN depletion. ARID1A wild-type cells exhibited significant G2/M phase arrest, whereas ARID1A-mutated cells showed diminished G2/M arrest and an increased proportion of cells in the G1 phase compared with the wild-type cells (Fig. 3d and Extended Data Fig. 5a). This effect was also observed in ARID1A-knockout cells (Extended Data Fig. 5b). Reintroducing ARID1A to ARID1A-mutated cells restored G2/M phase accumulation and reduced G1 phase arrest upon WRN depletion (Fig. 3e and Extended Data Fig. 5c). To determine whether the reduction in G2/M arrest in ARID1A-mutated cells resulted from changes in mitotic entry or exit, we examined p-H3 (Ser10), a marker of M phase, following paclitaxel treatment to block mitotic exit. WRN depletion reduced p-H3-positive cells in both ARID1A wild-type and ARID1A-mutated cells, indicating that the diminished G2/M arrest in ARID1A-mutated cells is not due to increased mitotic exit (Fig. 3f). Under paclitaxel treatment, WRN depletion induced G2 phase arrest in ARID1A wild-type cells, whereas ARID1A-mutated cells had a higher proportion of cells in the G1 phase (Fig. 3f and Extended Data Fig. 5d), confirming the differential cell cycle inhibitory effects associated with ARID1A mutation. Interestingly, despite slower proliferation, ARID1A-mutated cells displayed a slight but significant increase in the G2-to-M phase transition following *WRN* knockdown, as indicated by the proportion of p-H3-positive cells of total cells in the G2/M phase (Extended Data Fig. 5e). Consistently, G2/M arrest regulators (p-cdc2, Myt1, p-Wee1 and Wee1) were downregulated, whereas G1 arrest regulators (p21, p27, cyclin D1/D3, and CDK4/6) were upregulated in ARID1A-mutated cells (Fig. 3g and Extended Data Fig. 5f). These effects were also observed in ARID1A-knockout cells (Extended Data Fig. 5g) and were reversed upon ARID1A reintroduction (Fig. 3h and Extended Data Fig. 5h). These findings indicate that WRN depletion activates Chk1 and promotes G2/M arrest in ARID1A wild-type cells but primarily triggers Chk2 activation and G1 arrest in ARID1A-mutated cells.

To determine whether this distinct DNA damage response and cell cycle arrest pattern is also observed in other cell types, including MSS cells, we examined DNA damage signaling and cell cycle distribution following *WRN* knockdown. Microsatellite stable cells had only minor activation of DNA damage response upon *WRN* knockdown (Extended Data Fig. 6a). However, consistent with our earlier results, ARID1A wild-type MSI cells had predominantly Chk1 activation, whereas ARID1A-mutated MSI cells showed dominant Chk2 activation following *WRN* knockdown (Extended Data Fig. 6a). Similarly, upon WRN depletion, ARID1A wild-type MSI cells underwent significant G2/M arrest, while ARID1A-mutated MSI cells had an increased proportion of cells in G1 phase (Extended Data Fig. 6b). These results further support that WRN depletion differentially activates Chk signaling and induces distinct cell cycle arrest patterns based on ARID1A mutation status in MSI cancers.

### Chk1 blockade prompts compensatory Chk2 activation in ARID1A wild-type cells upon WRN depletion

ARID1A-deficient cancer cells have impaired ATR-Chk1 DNA damage response signaling due to reduced interaction between ARID1A and ATR. ^9, 27^ In addition, we found that basal expression and phosphorylation of ATR and Chk1 were lower in ARID1A-mutated cells, while Chk2 phosphorylation at baseline was elevated (Extended Data Fig. 6c). Based on these results, we hypothesized that the enhanced Chk2 activation observed in ARID1A-mutated cells upon WRN depletion is due to defective ATR-Chk1 signaling resulting from ARID1A loss-of-function mutation. To test this hypothesis, we investigated the effect of Chk1 inhibition on ARID1A wild-type cells following *WRN* knockdown. Chk1 inhibition with the Chk1 inhibitor prexasertib in WRN-depleted cells led to increased Chk2 phosphorylation and elevated DNA damage, as indicated by higher levels of phosphorylated p53 and γH2AX expression (Fig. 4a). Additionally, expression of G2/M checkpoint regulators (p-cdc2, Myt1, p-Wee1 and Wee1) was decreased, while expression of the G1 checkpoint regulator p21 was upregulated following *WRN* knockdown and prexasertib treatment (Fig. 4b). Consistent with these molecular changes, simultaneous inhibition of WRN and Chk1 reduced G2/M phase arrest and enhanced G1 phase arrest compared with WRN inhibition alone (Fig. 4c,d and Extended Data Fig. 6d). Furthermore, prexasertib treatment sensitized ARID1A wild-type cells to WRN depletion, as evidenced by decreased proliferation and increased apoptosis (Fig. 4e,f). These findings suggest that, under WRN depletion, Chk1 inhibition in ARID1A wild-type cells mimics the DNA damage response observed in ARID1A-mutated cells, in which compensatory Chk2 activation leads to increased G1 arrest. Thus, combining WRN depletion with Chk1 inhibition may be an effective strategy to target ARID1A wild-type cancer cells, potentially inducing vulnerabilities similar to those seen in ARID1A-mutated cells.

**Fig. 4.**
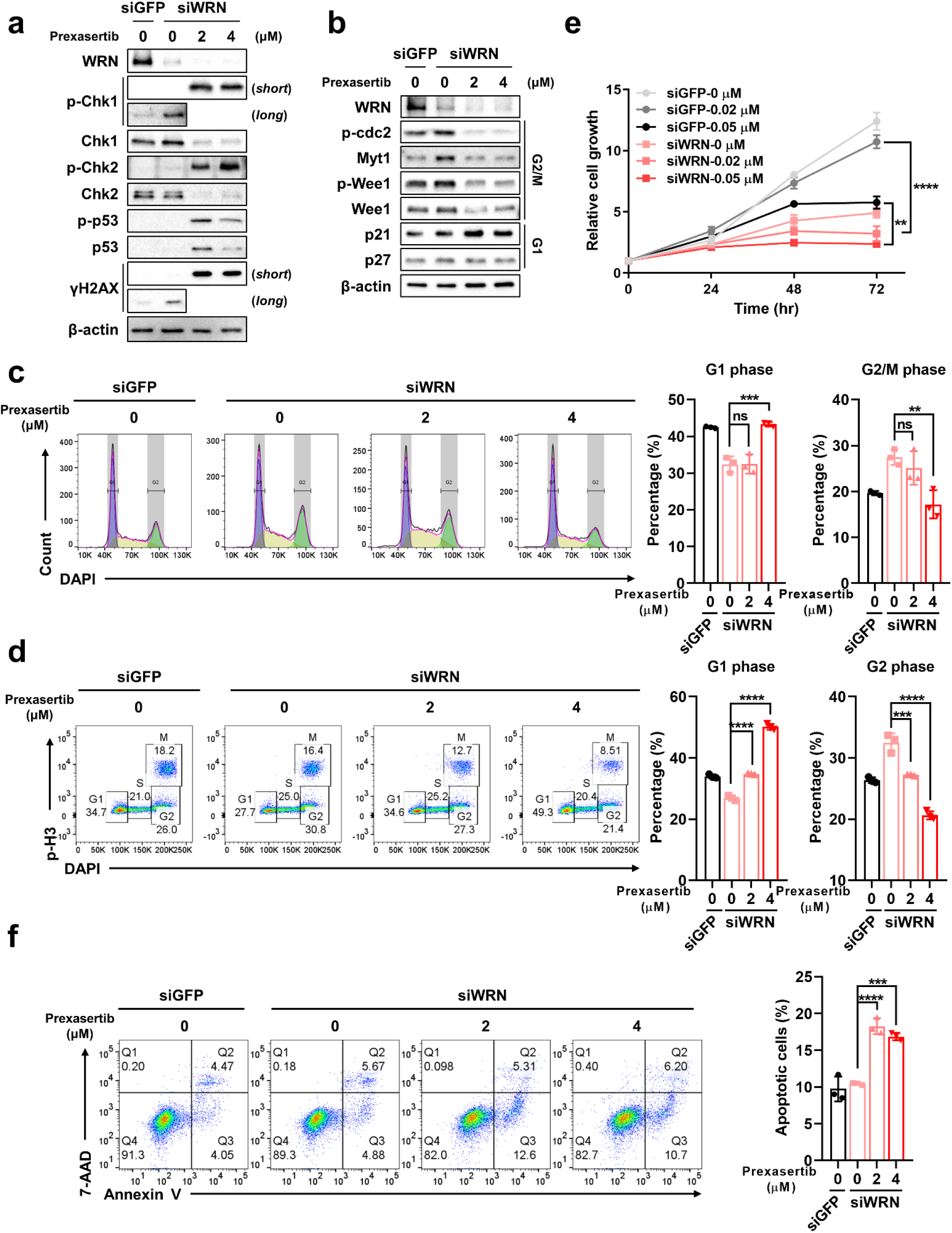
Chk1 inhibition activates Chk2 in ARID1A wild-type cells upon WRN depletion. (a) Western blot analysis of DNA damage response proteins in ARID1A wild-type cells upon *WRN* knockdown (6 hours), followed by prexasertib treatment (24 hours). (b) Western blot analysis of cell cycle regulatory proteins in ARID1A wild-type cells upon *WRN* knockdown (6 hours), followed by prexasertib treatment (24 hours). (c) G1 and G2/M phase proportions from cell cycle analysis in ARID1A wild-type cells upon *WRN* knockdown (6 hours), followed by prexasertib treatment (24 hours). Representative flow cytometry plots (left) and quantification (right) are shown. Data represent three independent experiments and are presented as mean ± standard deviation (SD). Statistical significance was determined using Tukey’s post hoc test following one-way analysis of variance (ANOVA). (d) Phospho-histone H3 staining assay in ARID1A wild-type cells upon *WRN* knockdown (6 hours), followed by prexasertib treatment (24 hours), showing G1 and G2 phase proportions. Cells were treated with paclitaxel (10 nM, 5 hours) at 24 hours after prexasertib treatment. Representative flow cytometry plots (left) and quantification (right) are shown. Data represent three independent experiments and are presented as mean ± SD. Statistical significance was determined using Tukey’s post hoc test following one-way ANOVA. (e) Cell proliferation measured using the WST-1 assay in ARID1A wild-type cells upon *WRN* knockdown (6 hours), followed by prexasertib treatment (24 hours), measured at 0, 24, 48, and 72 hours after transfection. Data represent three independent experiments and are presented as mean ± SD. Statistical significance was determined using Tukey’s post hoc test following one-way ANOVA. (f) Apoptosis assay in ARID1A wild-type cells upon *WRN* knockdown (6 hours), followed by prexasertib treatment (24 hours). Apoptosis was measured by annexin V/7-AAD staining 24 hours after prexasertib treatment. Representative flow cytometry plots (left) and quantification (right) are shown. Data represent three independent experiments and are presented as mean ± SD. Statistical significance was determined using Tukey’s post hoc test following one-way ANOVA. Statistical significance was defined as follows: not significant; ns, p > 0.05; *, p ≤ 0.05; **, p ≤ 0.01; ***, p ≤ 0.001; ****, p ≤ 0.0001.

### A WRN inhibitor suppresses cancer cell growth with distinct DNA damage signaling and cell cycle arrest in ARID1A-mutated cells

We next investigated the effect of pharmacological WRN inhibition using the selective WRN inhibitor NSC617145, which targets the WRN ATPase domain. ^28^ ARID1A-mutated cells exhibited increased sensitivity to NSC617145 compared with ARID1A wild-type cells, as evidenced by a lower IC_50_ in cytotoxicity assays and a concentration-dependent decline in colony formation (Fig. 5a,b). Similar cytotoxic effects were observed with other WRN inhibitors, NSC19630 and HRO761 (Extended Data Fig. 6e). Additionally, NSC617145 treatment led to a substantial increase in apoptosis in ARID1A-mutated cells (Fig. 5c). Consistent with *WRN* knockdown findings, Chk2 activation and elevated γH2AX levels were detected in ARID1A-mutated cells following NSC617145 treatment (Fig. 5d). Although NSC617145 induced a less pronounced G2/M phase accumulation compared with *WRN* siRNA treatment, ARID1A-mutated cells still had a higher proportion of cells in G1 phase and a lower proportion in G2/M phase compared with ARID1A wild-type cells (Fig. 5e). These findings were accompanied by differential expression of cell cycle regulators, mirroring the results observed with siRNA-mediated WRN depletion (Fig. 5f). These findings demonstrate that chemical inhibition of the ATPase activity of WRN exerts an anti-tumor effect in ARID1A-mutated cells.

**Fig. 5.**
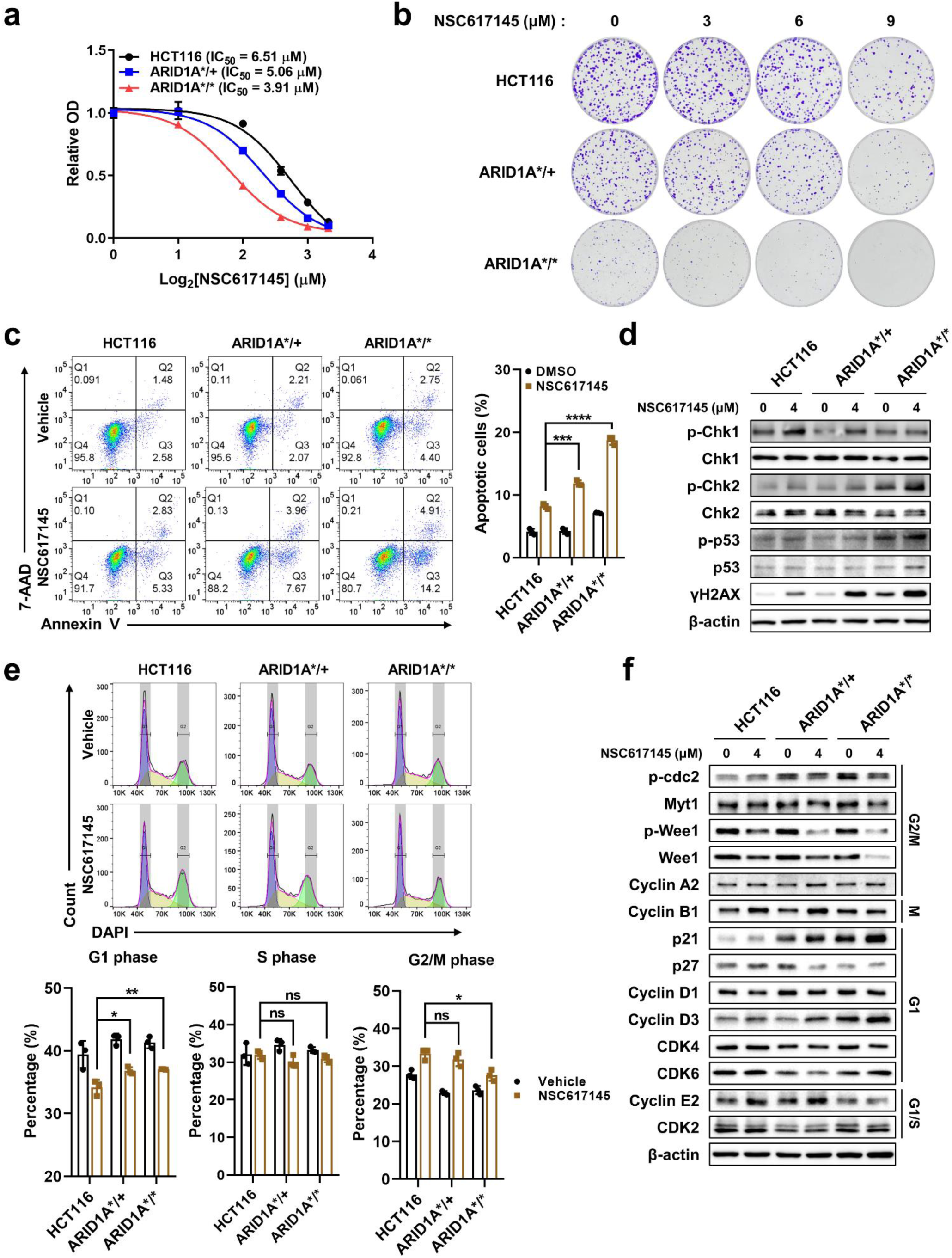
WRN inhibition by NSC617145 affects cell viability, apoptosis, and cell cycle regulation in ARID1A-mutated cells. (a) Drug cytotoxicity assay in ARID1A wild-type and ARID1A-mutated cells with NSC617145 treatment (72 hours). Drug concentrations ranged from 0 to 10 μM. IC_50_ values are indicated on the plot with a fitted dose-response curve. (b) Representative images of the colony formation assay in ARID1A wild-type and ARID1A-mutated cells with NSC617145 treatment, assessed after 10 days of culture. (c) Apoptosis assay in ARID1A wild-type and ARID1A-mutated cells with NSC617145 treatment (8 μM), measured by annexin V/7-AAD staining at 48 hours after NSC617145 treatment. Representative flow cytometry plots (left) and quantification (right) are shown. Data represent three independent experiments and are presented as mean ± standard deviation (SD). Statistical significance was determined using Tukey’s post hoc test following one-way analysis of variance (ANOVA). (d) Western blot analysis of DNA damage response proteins in ARID1A wild-type and ARID1A-mutated cells with NSC617145 treatment (4 μM, 24 hours). (e) Cell cycle analysis in ARID1A wild-type and ARID1A-mutated cells with NSC617145 treatment (4 μM, 24 hours). Representative flow cytometry plots (top) and quantification (bottom) are shown. Data represent three independent experiments and are presented as mean ± SD. Statistical significance was determined using Tukey’s post hoc test following one-way ANOVA. (f) Western blot analysis of cell cycle regulatory proteins in ARID1A wild-type and ARID1A-mutated cells with NSC617145 treatment (4 μM, 24 hours). Statistical significance was defined as follows: not significant; ns, p > 0.05; *, p ≤ 0.05; **, p ≤ 0.01; ***, p ≤ 0.001; ****, p ≤ 0.0001.

### WRN depletion in ARID1A-mutated cells suppresses tumor growth in a xenograft model

To evaluate the effect of WRN depletion in vivo, we conducted tumor growth assays in NOD/SCID/IL-2γ-receptor-null (NSG) mice. Prior to the experiment, we confirmed that ARID1A wild-type and ARID1A-mutant cells had similar growth rates in vivo, which contrasted with the differential rates observed in vitro (Extended Data Fig. 7a). We used a doxycycline-inducible shRNA system to deplete WRN, using shWRN-C911 (non-targeting shRNA) as a negative control. ^13^ Doxycycline-induced WRN depletion using the shRNA for *WRN* (shWRN1 and shWRN2) significantly inhibited tumor growth in ARID1A-mutated cells, whereas the inhibition of WRN in ARID1A wild-type cells was less pronounced (Fig. 6a-d). As expected, the shWRN-C911s control had little effect on tumor growth regardless of doxycycline treatment (Extended Data Fig. 7b-e). Immunohistochemistry analysis confirmed reduced WRN expression following doxycycline treatment, which was accompanied by decreased Ki-67 expression and increased p21 and γH2AX levels in ARID1A-mutant tumors (Fig. 6e,f and Extended Data Fig. 7f).

**Fig. 6.**
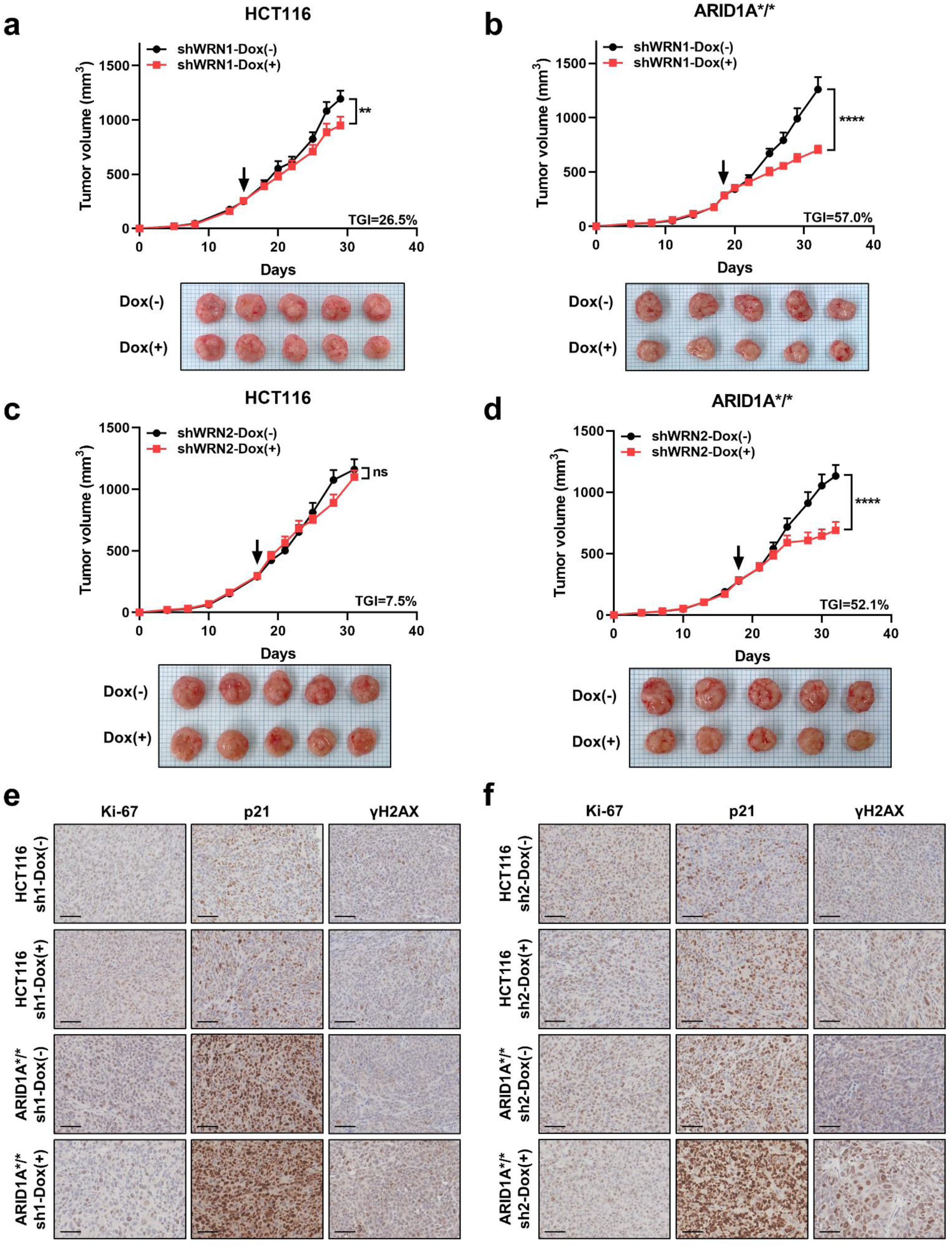
WRN depletion suppresses tumor growth in ARID1A-mutated cells in vivo. (a,b) Tumor growth curves for shWRN1 xenografts with ARID1A wild-type HCT116 (a) and ARID1A-mutated HCT116 cells (b) in NSG mice (top, n = 5 per group). The arrow indicates the start of doxycycline (DOX) treatment. Representative tumor images from sacrificed mice at the end of the experiment are shown (bottom). Data are presented as mean ± standard error of the mean (SEM). Statistical significance was determined using Sidak’s post hoc test following two-way analysis of variance (ANOVA). (c,d) Tumor growth curves for shWRN2 xenografts with ARID1A wild-type HCT116 (c) and ARID1A-mutated HCT116 cells (d) in NSG mice (top, n = 5 per group). The arrow indicates the start of DOX treatment. Representative tumor images from mice killed at the end of the experiment are shown (bottom). Data are presented as mean ± SEM. Statistical significance was determined using Sidak’s post hoc test following two-way ANOVA. (e,f) Representative immunohistochemistry images of Ki-67, p21, and γH2AX staining in tumors from shWRN1 (e) and shWRN2 (f) xenografts. Scale bar, 50 μm. Statistical significance was defined as follows: not significant; ns, p > 0.05; *, p ≤ 0.05; **, p ≤ 0.01; ***, p ≤ 0.001; ****, p ≤ 0.0001. TGI, tumor growth inhibition.

To further assess the therapeutic potential of WRN inhibition using a WRN inhibitor in vivo, we tested NSC617145 in mice implanted with ARID1A wild-type or ARID1A-mutated HCT116 cells. NSC617145 treatment reduced tumor growth of ARID1A-mutated cells, although its anti-tumor effect was less pronounced compared with WRN depletion via shRNA (Extended Data Fig. 8a,b). Overall, these findings suggest that WRN inhibition selectively suppresses tumor growth of ARID1A-mutated cells in vivo, supporting WRN as a potential therapeutic target in ARID1A-mutated cancers.

### Additional p21 inhibition in WRN-depleted conditions induces mitotic catastrophe selectively in ARID1A-mutated cells

We demonstrated that Chk1 defects in ARID1A-mutated cells result in G1 arrest upon WRN depletion, whereas Chk1 activation leads to G2 arrest in ARID1A wild-type cells. Based on these findings, we hypothesized that G2 arrest is also impaired in ARID1A-mutated cells upon WRN depletion. Consequently, we proposed that escaping G1 arrest in ARID1A-mutated cells under WRN depletion could lead to abnormal cell cycle progression, ultimately resulting in mitotic catastrophe, a form of cell death caused by improper cell cycle progression in the presence of DNA damage, such as premature mitotic entry. ^29^ This mechanism would selectively target ARID1A-mutated cells, as ARID1A wild-type cells could still arrest in the G2 phase even if they bypass G1 arrest.

p21 inhibition can facilitate escape from G1 arrest, ^30^ we combined WRN depletion with *p21* knockdown. As hypothesized, this combination selectively induced apoptosis in ARID1A-mutated cells (Fig. 7a,b). Cell cycle analysis revealed that concurrent WRN depletion and *p21* knockdown reduced G1 arrest as expected, but the proportion of cells in the G2/M phase was decreased in ARID1A-mutated cells (Fig. 7c Extended Data Fig. 9a). Instead, there was a notable increase in the sub-G1 population, a marker of apoptotic cells (Extended Data Fig. 9a). These results suggest that escaping G1 arrest drives cell cycle progression despite DNA damage, leading to mitotic catastrophe. To further assess cell cycle progression, we conducted a p-H3 staining assay following paclitaxel treatment. Concurrent knockdown of *WRN* and *p21* resulted in increased M phase accumulation without altering the proportion of G2 phase cells compared with *WRN* knockdown alone in ARID1A-mutated cells (Fig. 7d and Extended Data Fig. 9b). Additionally, under these conditions, the ratio of G2 to M phase progression in ARID1A-mutated cells was higher than in ARID1A wild-type cells (Fig. 7d). Thus, p21 inhibition facilitates the escape from G1 arrest, and combined inhibition of p21 and WRN selectively induces cell death by promoting abnormal mitotic progression in ARID1A-mutated cells. To validate this concept pharmacologically, we used UC2288, a p21 inhibitor, in combination with NSC617145, which demonstrated a synergistic effect, significantly increasing apoptosis (Fig. 7e,f). Furthermore, immunofluorescence staining confirmed mitotic catastrophe, as indicated by abnormal nuclear morphology, which was significantly increased in ARID1A-mutated cells following *p21* and *WRN* knockdown (Fig. 7g). Therefore, p21 inhibition enhances the cytotoxicity of WRN inhibition in ARID1A-mutated cells, providing a combinatorial therapeutic strategy for ARID1A-mutant cancers.

**Fig. 7.**
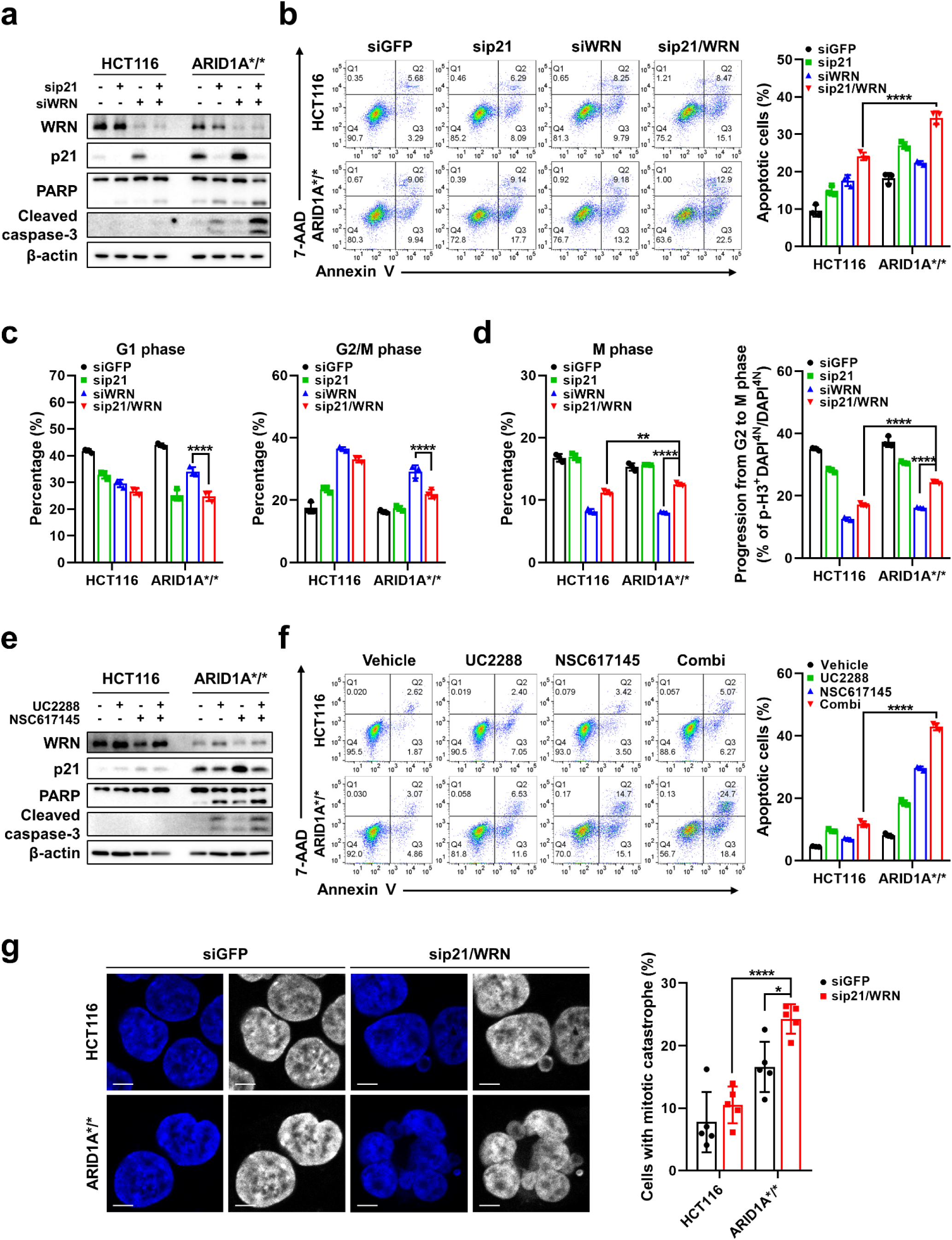
Dual inhibition of WRN and p21 induces apoptosis and mitotic catastrophe in ARID1A-mutated cells. (a) Western blot analysis of apoptosis markers following *WRN* and *p21* knockdown in ARID1A wild-type and ARID1A-mutated cells. (b) Apoptosis assay following *WRN* and *p21* knockdown in ARID1A wild-type and ARID1A-mutated cells, measured by annexin V/7-AAD staining at 48 hours after transfection. Representative flow cytometry plots (left) and quantification (right) are shown. Data represent three independent experiments and are presented as mean ± standard deviation (SD). Statistical significance was determined using Dunnett’s post hoc test following one-way analysis of variance (ANOVA). (c) G1 and G2/M phase proportions from cell cycle analysis following *WRN* and *p21* knockdown in ARID1A wild-type and ARID1A-mutated cells. Data represent three independent experiments and are presented as mean ± SD. Statistical significance was determined using Dunnett’s post hoc test following one-way ANOVA. (d) M phase proportion and G2-to-M phase progression from phospho-histone H3 (p-H3) staining assay following *WRN* and *p21* knockdown in ARID1A wild-type and ARID1A-mutated cells. Cells were treated with paclitaxel (10 nM, 5 hours) at 48 hours after transfection. The proportion of p-H3-positive cells among the total G2/M phase is considered as cells progress from G2 to M phase. Data represent three independent experiments and are presented as mean ± SD. Statistical significance was determined using Dunnett’s post hoc test following one-way ANOVA. (e) Western blot analysis of apoptosis markers following NSC617145 (4 μM) and UC2288 (5 μM) treatment for 24 hours in ARID1A wild-type and ARID1A-mutated cells. (f) Apoptosis assay following NSC617145 (4 μM) and UC2288 (5 μM) treatment in ARID1A wild-type and ARID1A-mutated cells, measured by annexin V/7-AAD staining at 48 hours after drug treatment. Representative flow cytometry plots (left) and quantification (right) are shown. Data represent three independent experiments and are presented as mean ± SD. Statistical significance was determined using Dunnett’s post hoc test following one-way ANOVA. (g) Immunofluorescence images (left) of nuclei stained with DAPI following *WRN* and *p21* knockdown in ARID1A wild-type and ARID1A-mutated cells. Quantification (right) shows the percentage of mitotic catastrophe-positive cells. Cells undergoing mitotic catastrophe were quantified from multiple independent images (n = 5). Scale bar, 5 μm. Statistical significance was defined as follows: not significant; ns, p > 0.05; *, p ≤ 0.05; **, p ≤ 0.01; ***, p ≤ 0.001; ****, p ≤ 0.0001.

As p53 is an upstream regulator of p21 and plays a role in cell cycle regulation, including G1 arrest, ^31, 32^ we performed *p53* knockdown to determine whether inhibiting p53 would replicate the effects of p21 inhibition. However, *p53* knockdown in combination with WRN depletion did not enhance apoptosis in ARID1A-mutated cells (Extended Data Fig. 10a,b). Similar to the results for *p21* knockdown, cell cycle analysis revealed that *p53* knockdown facilitated escape from G1 arrest (Extended Data Fig. 10c). However, unlike p21 inhibition, *p53* knockdown increased G2/M phase accumulation under WRN depletion in ARID1A-mutated cells (Extended Data Fig. 10c). Moreover, p-H3 staining following paclitaxel treatment showed that *p53* knockdown led to G2 phase accumulation without promoting M phase progression (Extended Data Fig. 10d). When we examined the expression of G2/M regulators, the expression of Wee1 and phosphorylation of cdc2, Wee1, and Chk1 were significantly enhanced by the depletion of p53 combined with *WRN* knockdown (Extended Data Fig. 10e). Additionally, cyclin B1, a key regulator of mitotic entry via nuclear localization, ^33^ was also upregulated in p53-inhibited cells (Extended Data Fig. 10e); however, immunofluorescence analysis revealed its retention in the cytoplasm (Extended Data Fig. 9c), indicating that the increased cyclin B1 was not functionally active for facilitating mitotic entry. These results suggest that in contrast to p21 inhibition, p53 inhibition activates G2 arrest upon WRN depletion, thereby preventing mitotic catastrophe while permitting G1 progression.

In an in vivo cell line xenograft mouse model, the combination of NSC617145 and UC2288 significantly reduced tumor growth in ARID1A-mutated HCT116 cell xenografts (Fig. 8a). Furthermore, colon cancer patient-derived xenograft (PDX) models harboring an ARID1A mutation (S764fs) also had reduced tumor growth following the combination treatment (Fig. 8b). Although the NSC617145 monotherapy and the combination treatment led to a notable 15% to 20% reduction in mouse body weight, this change remained within a tolerable range (Extended Data Fig. 8c,d). Immunohistochemistry analysis of PDX tissues from treated mice showed decreased Ki-67 and increased cleaved caspase-3, γH2AX, and p21 in the combinatorial treatment group (Fig. 8c). These findings indicate that simultaneous inhibition of WRN and p21 represents a promising combination treatment strategy for ARID1A-mutated cancers.

**Fig. 8.**
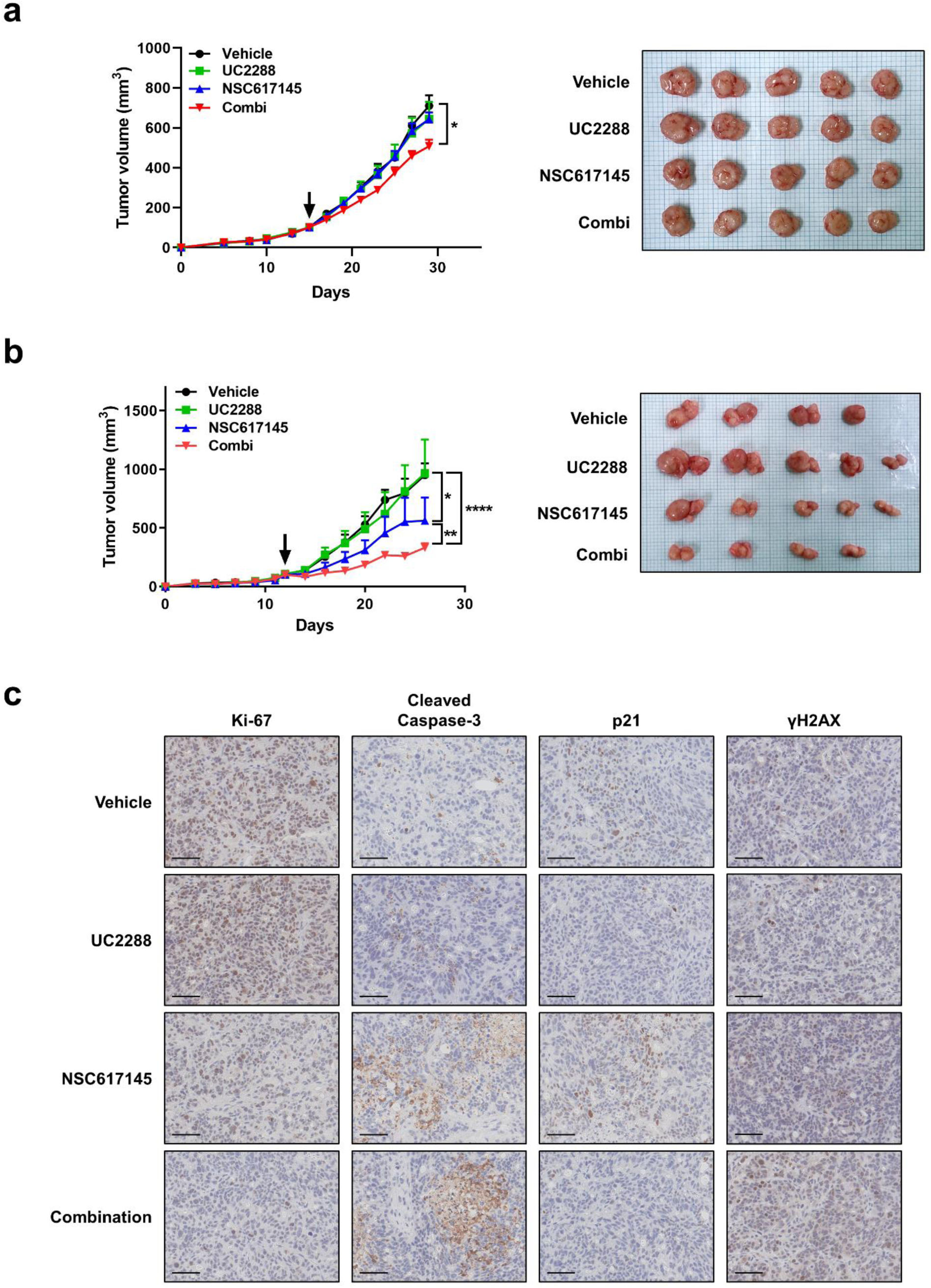
Combinatorial inhibition of WRN and p21 suppresses tumor growth in ARID1A-deficient tumors in vivo. (a,b) Tumor growth curves following combinatorial treatment with NSC617145 (20 mg/kg) and UC2288 (30 mg/kg) in ARID1A-mutated HCT116 cells as xenograft models (a) and patient-derived xenograft models (b) from patients with ARID1A mutation and colon cancer (left, n = 5 per group). The arrow indicates the start of drug treatment. Representative tumor images from sacrificed mice at the end of the experiment are shown (right). Data are presented as mean ± standard error of the mean (SEM). Statistical significance was determined using Sidak’s post hoc test following two-way analysis of variance. (c) Representative immunohistochemistry images of PDX tumor samples stained for Ki-67, cleaved caspase-3, p21, and γH2AX. Scale bar, 50 μm. Statistical significance was defined as follows: not significant; ns, p > 0.05; *, p ≤ 0.05; **, p ≤ 0.01; ***, p ≤ 0.001; ****, p ≤ 0.0001.

## Discussion

In this study, we explored new therapeutic strategies targeting ARID1A-mutated cancers, focusing on synthetic lethal interactions and vulnerabilities linked to DNA damage response and cell cycle regulation. In ARID1A-mutated cells, targeting the DNA helicase WRN triggered distinct ATR-Chk1 activation, leading to G1 arrest, and further targeting of p21 alongside WRN inhibition induced mitotic catastrophe-mediated apoptosis. Our study expands on the landscape of targeted therapeutics in ARID1A-mutated cancers.

Previous studies have identified various synthetic lethal targets in ARID1A-mutated cancers, including cell cycle regulators and DNA damage response components (e.g., PARP, ATR, and AURKA-CDC25C), ^9, 10, 12^ epigenetic regulators (e.g., EZH2 and HDAC6), ^8, 11^ chromatin remodeling components that function as a compensatory partner (e.g., ARID1B), ^7^ and crucial signaling pathways such as PI3K/AKT (e.g., PIK3CA). ^34^ Our analysis demonstrated that genes with differential dependency in ARID1A-mutated cells were enriched in pathways associated with DNA damage response and cell cycle regulation, underscoring the critical role of these processes in the viability of ARID1A-mutated cancers. These results suggest that targeting these pathways could offer a selective therapeutic strategy for treating ARID1A-mutated cancers.

We propose WRN as a promising therapeutic target in ARID1A-mutated cancers, particularly within the context of MSI. While WRN has already been identified as a synthetic lethal target in MSI cancers, our study reveals that ARID1A mutations are an additional and crucial determinant of *WRN* dependency. We suggest that ARID1A mutations could serve as a biomarker for predicting the efficacy of WRN inhibition therapies in patients with MSI cancers. Given that ARID1A mutations frequently occur in MSI-high colorectal cancers, ^15, 16^ the relationship among ARID1A, WRN, and MSI presents an important therapeutic opportunity. Furthermore, although our study focused on colorectal cancer cell lines, MSI status is a characteristic of several cancer types, suggesting that our findings may be applicable to a broad range of MSI-associated cancers.

Our investigation into *WRN* dependency uncovered that WRN depletion activates distinct DNA damage signaling pathways depending on the ARID1A mutation status. Specifically, WRN depletion activated Chk1-mediated DNA damage responses in ARID1A wild-type cells, while Chk2-mediated responses were predominant in ARID1A-mutated cells. These differences resulted in unique cell cycle arrest outcomes, with Chk1-driven G2 arrest in ARID1A wild-type cells and Chk2-driven G1 arrest in ARID1A-mutated cells. These findings highlight the impact of ARID1A mutations on DNA damage response, indicating that other DNA damage-based therapies could also be effective against ARID1A-mutated cancers.

We further investigated a compensatory DNA damage response mechanism in ARID1A-mutated cancers following WRN depletion. ARID1A directly interacts with ATR and is essential for ATR activation in response to ionizing radiation-induced DNA damage. ^9^ Consistently, our data show that basal expression and activation of the ATR-Chk1 axis are downregulated in ARID1A-mutated cells (Extended Data Fig. 6c). This impairment leads to compensatory activation of the Chk2 pathway upon WRN depletion. These findings suggest a new therapeutic approach: combining Chk1 and WRN inhibition in ARID1A wild-type cancers to mimic the apoptotic response seen in ARID1A-mutated cancers under WRN inhibition.

In in vivo cell line xenograft mouse models, genetic inhibition of *WRN* demonstrated substantial anti-tumor effects, reducing tumor growth by approximately 50% in ARID1A-mutated cells, while having only minimal impact on ARID1A wild-type tumors. This finding reinforces WRN as a selective and effective synthetic lethal target in ARID1A-mutated cancers. However, pharmacological inhibition using NSC617145 was less effective compared with the genetic knockdown of *WRN*. This effect is probably due to the limited efficacy and poor bioavailability of the WRN inhibitor NSC617145, which may hinder its ability to effectively target and inhibit WRN in vivo, reducing its therapeutic potential. Therefore, the emergence of new and more specific WRN inhibitors, such as HRO761, ^35^ may offer greater efficacy for targeting ARID1A-mutated cancers.

Additionally, we propose a new therapeutic strategy that combines WRN inhibition with p21 inhibition in ARID1A-mutated cancers. This approach leverages the Chk1-G2 arrest defects in ARID1A-mutated cells, enhancing cytotoxicity by inducing mitotic catastrophe. By disrupting G1 arrest through p21 inhibition, we forced cell cycle progression in the presence of DNA damage, effectively inducing mitotic catastrophe. This strategy represents a promising avenue for treating ARID1A-mutated cancers and highlights the potential of targeting cell cycle progression defects in ARID1A-mutated cancers.

Overall, our study provides comprehensive insights into the vulnerabilities of ARID1A-mutated cancers, revealing WRN as a potent therapeutic target and introducing innovative strategies to exploit cell cycle dysregulation. These findings have significant implications for developing targeted treatments for ARID1A-mutated MSI-high cancers, paving the way for precision oncology approaches that leverage the unique genetic and regulatory landscapes of these tumors.

## Materials and Methods

### Differential genetic dependency analysis

We obtained CRISPR gene-dependency data from the DepMap Public 22Q2 dataset (Project Achilles). Gene-dependency scores for each cell line in Cancer Cell Line Encyclopedia (CCLE) were estimated using the CERES algorithm, which adjusts for confounding effects such as gene copy number variations. ^18^ Mutation profiles of cancer cell lines were curated from the Cancer Cell Line Encyclopedia mutation dataset provided by DepMap, incorporating hotspot mutations reported in The Cancer Genome Atlas (TCGA) and COSMIC databases (e.g., *PIK3CA*, *BRAF*, *KRAS*). For tumor suppressor genes (e.g., *ARID1A*, *PTEN*, *TP53*, *FBXW7*, *ATM*, *APC*), predicted deleterious mutations were also considered, as loss-of-function alterations in these genes are functionally relevant. ^36–41^ Cell lines harboring mutations with functional alterations were classified as mutant, whereas cell lines without any mutations, including non-hotspot mutations, were classified as wild-type. The MSI status for each cell line was obtained from phase II of the CCLE project. ^42^

Differential analysis of gene-dependency scores between ARID1A wild-type (n = 901) and ARID1A-mutant (n = 115) cell lines was conducted using the limma package in R. We fitted a linear model for each gene, comparing the dependency scores between the wild-type and mutant groups. Empirical Bayes moderation was applied to calculate p-values, and multiple hypothesis testing was controlled using Benjamini–Hochberg false discovery rate correction.

### ClueGO enrichment analysis

Functional enrichment analysis of differential dependency genes (n = 50) was performed using the ClueGO plugin (version 2.5.10) in Cytoscape software (version 3.10.2). For both Gene Ontology Biological Process and REACTOME Pathway gene sets, the following settings were applied: evidence codes, all; network specificity, medium; kappa score threshold, 0.4; minimum number of genes, 2; minimum percentage of genes, 4%.

### Gene set enrichment analysis

Gene expression data from TCGA were obtained via cBioPortal (https://www.cbioportal.org). Raw data were normalized and preprocessed using reads per kilobase of transcript per million mapped reads. Gene set enrichment analysis (GSEA) was performed using the GSEA software (Broad Institute, version 4.1.0) by comparing groups stratified by the median WRN expression level. The analysis was conducted using hallmark gene sets from the Molecular Signatures Database (version 2024.1).

### Correlation analysis

Gene expression data from The Cancer Genome Atlas and the Cancer Cell Line Encyclopedia were obtained via cBioPortal (https://www.cbioportal.org). The dataset included 592 samples from TCGA and 1,156 samples from the CCLE, including 59 large-intestine and small-intestine cancer cell lines. Pearson correlation analysis was conducted using R (version 4.4.2).

### Cell culture

We purchased human colon cancer HCT116 isogenic cell lines with wild-type ARID1A, with heterozygously mutated ARID1A (Q456*/+, HD 104-031), or with homozygously mutated ARID1A (Q456*/Q456*, HD 104-049) from Horizon Discovery. The cells were cultured in RPMI 1640 medium (Cytiva, SH30027.01) supplemented with 10% fetal bovine serum (Cytiva, SH30919.03) and 1% penicillin-streptomycin (Gibco, 15140-122). Cells were incubated in a humidified incubator at 37°C with 5% CO_2_.

### Quantitative PCR

Total RNA was extracted using the RNeasy Mini Kit (QIAGEN, 74106) in accordance with the manufacturer’s instructions. Reverse transcription was performed using the 5× PrimeScript RT Master Mix (Takara Bio, RR036A) following the provided protocol. qPCR was performed using iQ SYBR Green Supermix (Bio-Rad Laboratories, 1708882) on a Bio-Rad Laboratories CFX96 real-time PCR system. Relative gene expression was calculated using the 2^−ΔΔCt^ method, with *GAPDH* as the reference gene. Primer sequences are provided in Supplementary Information 1.

### Immunoblotting

Cellular proteins were extracted in RIPA buffer (Thermo Fisher Scientific, 89900) supplemented with complete protease inhibitor cocktail (Roche, 11873580001) and phosphatase inhibitor cocktail (Roche, 04906837001) for 30 minutes at 4°C. Proteins were separated by SDS-PAGE and transferred to nitrocellulose membranes (GE Healthcare, 10600002). The membranes were blocked with 5% (w/v) skim milk powder (BD Biosciences, 232100) in Tris-buffered saline with 0.1% Tween 20 (TBS-T; Biosesang, TR2007-100-74) at room temperature for 1 hour and then incubated with primary antibodies overnight at 4°C. After washing with TBS-T, membranes were incubated with horseradish peroxidase-conjugated secondary antibodies: anti-rabbit IgG (Invitrogen, 31460) or anti-mouse IgG (Invitrogen, 31430). Bound antibodies were detected by enhanced chemiluminescence (GE Healthcare) in accordance with the manufacturer’s instructions. Band intensity quantification was performed using ImageJ software (National Institutes of Health). Antibody details and dilution ratios are provided in Supplementary Information 2.

### Nascent RNA synthesis and quantification

Nascent RNA synthesis was assessed using the Click-iT Nascent RNA Capture Kit (Invitrogen, C10365) in accordance with the manufacturer’s instructions. Cells were incubated with 0.5 mM 5-ethynyl uridine (EU) for 1 hour. Total RNA, including EU-labeled RNA, was extracted using TRIzol Reagent (Invitrogen, 15596018). A copper-catalyzed click reaction was performed between EU-labeled RNA and biotin azide, and then biotinylated nascent RNA was captured using streptavidin magnetic beads. The captured nascent RNA was reverse transcribed into cDNA, and qPCR was performed. The percent input for each sample was calculated using this formula: % input = 100 × 2^−(Ct[Nascent^ ^RNA]−Ct[Adjusted^ ^Input])^. The qPCR primers for *WRN* were as follows: ^43^ forward, 5′-TGCTAGTGATTGCTCTTTCCTG-3′; reverse, 5′-CTTTGCCAAGTTTCCCTCTATTG-3′.

### Transfection

Knockdown of genes was achieved by siRNA transfection conducted using Lipofectamine 2000 (Invitrogen, 11668019) in accordance with the manufacturer’s instructions. *GFP* siRNA (sc-45924), *WRN* siRNA (sc-36843), *p21* siRNA (sc-29427), and *p53* siRNA (sc-29435) were purchased from Santa Cruz Biotechnology. *GFP* siRNA was used as a negative control. Knockdown was confirmed by immunoblot 2 days after transfection.

### Chromatin immunoprecipitation followed by qPCR (ChIP-qPCR)

Chromatin immunoprecipitation was performed using the SimpleChIP Enzymatic Chromatin IP Kit (Cell Signaling Technology, 9003) in accordance with the manufacturer’s instructions. Cells were crosslinked with 1% paraformaldehyde for 10 minutes and the reaction was quenched with glycine. Chromatin was fragmented using micrococcal nuclease (MNase) digestion, and nuclei were further disrupted by sonication. For each sample, 10 μg of chromatin was immunoprecipitated with either anti-ARID1A (1:100; Cell Signaling Technology, 12354) or anti-RNAPII subunit B1 (1:50; Cell Signaling Technology, 13523). The chromatin and antibody mixtures were incubated overnight at 4°C, and the chromatin fragments were eluted. Samples were reverse crosslinked with proteinase K at 65°C for 2 hours, and DNA was purified using the ChIP DNA Clean & Concentrator (Zymo Research, D5205). Quantitative PCR was performed using the following formula to calculate percent input: % input = 100 × 2^−(Ct[IP] − Ct[Adjusted Input])^. The *WRN* promoter primers used for ChIP-qPCR were as follows: forward, 5′-CGGGTCGGGTAGGTCT-3′; reverse, 5′-CAGGCTAGACACCAAATCAGC-3′.

### WST-1 assay

Cells were cultured after transfection with siRNA or after drug treatment in a 96-well plate. The cells were treated with water-soluble tetrazolium salt (WST) substrate (DoGenBio, EZ-1000) for 1 hour in a humidified incubator at 37°C with 5% CO_2_ and then analyzed on a microplate reader (Epoch) at 450 nm. An equivalent volume of RPMI 1640 medium with control siRNA or vehicle was used as a background control.

### BrdU-incorporation assay

Cell proliferation was assessed using a colorimetric 5-bromo-2′-deoxyuridine (BrdU) cell proliferation kit (Roche, 11647229001) in accordance with the manufacturer’s instructions. Briefly, transfected cells were cultured for 24 hours and labeled with BrdU for 8 hours. Then, the cells were fixed with FixDent for 30 minutes and incubated with anti-BrdU-peroxidase for 90 minutes. After three washes with washing solution, the substrate solution was added for 3 to 5 minutes in the dark, and the absorbance of the samples was measured on a microplate reader (Epoch) at 450 nm.

### Colony formation assay

Cells were cultured in 60-mm dishes or 6-well plates after transfection with siRNA or after drug treatment. Cell medium was replaced every 2 to 3 days. Colonies were fixed with cold 100% methanol for 30 minutes and stained with 1% crystal violet (Sigma-Aldrich, V5265) at room temperature for 10 minutes.

### Apoptosis assay

Cells were harvested with cold Dulbecco’s phosphate-buffered saline (DPBS; Welgene, LB001-02) and stained with FITC annexin Ⅴ (BD Biosciences, 556420) and 7-amino-actinomycin D (7-AAD; BD Biosciences, 559925) in 1× Annexin Ⅴ Binding Buffer (BD Biosciences, 556454) for 15 minutes at room temperature. The annexin Ⅴ-positive and 7-AAD-positive cells were detected with an LSRFortessa X-20 flow cytometer (BD Biosciences).

### Comet assay

DNA damage was assessed using the CometAssay Single Cell Gel Electrophoresis Assay (Trevigen, #4250-050-K) in accordance with the manufacturer’s instructions. Cells suspended in DPBS were mixed with LMAgarose at a 1:10 (v/v) ratio and spread onto CometSlides. After incubating for 30 minutes at 4°C in the dark, cells were lysed with lysis buffer overnight at 4°C. For neutral comet assay, lysed cells were electrophoresed at 21 volts for 45 minutes at 4°C. For alkaline comet assay, lysed cells were incubated in alkaline unwinding solution for 1 hour at 4°C and electrophoresed at 21 volts for 30 minutes at 4°C. Following electrophoresis, samples were washed with 70% ethanol and dried at 37°C for 10 minutes. Slides were mounted with fluorescence mounting medium (Agilent, S3023) and imaged using an LSM 980 confocal microscope (ZEISS).

### Immunofluorescence staining

Cells were grown on coverslips coated with poly-L-lysine (Sigma-Aldrich, P4832). Cells were fixed with 4% paraformaldehyde (Biosesang, PC2031-100-00) and permeabilized with 0.25% Triton X-100 (Sigma-Aldrich, T8787). Cells were blocked with 5% bovine serum albumin in DPBS at 37°C for 1 hour and incubated with primary antibodies at 4°C overnight. Samples were incubated with fluorescently labeled secondary antibodies at room temperature for 1 hour. After washing with DPBS, samples were mounted using 4’,6-diamidino-2-phenylindole (DAPI)-containing mounting medium (Sigma-Aldrich, DUO82040) at room temperature for 30 minutes and sealed with nail polish. Slides were imaged using an LSM 980 confocal microscope (ZEISS). Immunofluorescent foci were quantified using ImageJ software (National Institutes of Health).

### Cell cycle analysis

Cells were fixed in cold 70% ethanol and incubated with 0.1% (v/v) Triton X-100 (Sigma-Aldrich, T8787) and 1 μg/mL DAPI (Sigma-Aldrich, D9542) at room temperature for 30 minutes. The DNA content was analyzed with an LSRFortessa flow cytometer (BD Biosciences) using FACSDiva software (BD Biosciences).

### Phospho-histone H3 (p-H3) staining assay

Cells were fixed in cold 70% ethanol and permeabilized with 0.25% (v/v) Triton X-100 (Sigma-Aldrich, T8787) in DPBS at room temperature for 15 minutes. The cells were incubated with anti-phospho-histone H3 (anti-p-H3) antibody conjugated to Alexa Fluor 647 fluorescent dye (Cell Signaling Technology, 3458) at 4°C overnight. Subsequently, samples were stained with 1 μg/mL DAPI at room temperature for 30 minutes and analyzed on an LSRFortessa flow cytometer (BD Biosciences) using FACSDiva software (BD Biosciences).

### Generation of cell lines with ARID1A-knockout or overexpression

ARID1A-knockout constructs, including lentiCRISPR v2 (plasmid 52961), psPAX2 (plasmid 12260), and pCMV-VSV-G (plasmid 8454) were purchased from Addgene. The sgRNAs for *ARID1A* are as follows: sgARID1A-1, 5′-CGGTACCCGATGACCATGC-3′; sgARID1A-2, 5′-ATGGTCATCGGGTACCGCTG-3′. The sgRNAs were introduced to the lentiCRISPR v2 vector using Golden Gate assembly. The lentiviral plasmids were transfected into 293FT cells (Invitrogen, R70007) with Lipofectamine 2000 (Invitrogen, 11668019). Two days after transfection, the supernatant was collected and lentiviruses were concentrated using a Lenti-X Concentrator (Takara Bio, 631232). HCT116 cells were transduced with concentrated lentivirus and ARID1A-knockout clones were selected with 1 μg/mL puromycin. The *ARID1A* knockout was verified by immunoblot.

The ARID1A-overexpression vector pLenti-puro-ARID1A (plasmid 39478) was purchased from Addgene. The lentiviral production was performed as described above and concentrated lentivirus was added to ARID1A-mutated (Q456*/Q456*) cells. An ARID1A-overexpression clone was selected by verifying by immunoblot.

### In vivo mouse xenografts

All animal studies were approved by the Institutional Animal Care and Use Committee of Seoul National University (SNU-240205-1-2). Six- to eight-week-old NOD/SCID/IL-2γ-receptor-null (NSG) mice were obtained from The Jackson Laboratory for xenograft experiments.

To study the effects of *WRN* knockdown using shRNA, we subcutaneously implanted 2 × 10^6^ cells of the following cell types into the right flank of NSG mice: ARID1A wild-type or ARID1A-mutant (Q456*/Q456*) HCT116 cells transduced with doxycycline-inducible *WRN* shRNA1/2 or *WRN*-C911 shRNA1/2 plasmids (Addgene, plasmids 125783, 125784, 125785, 125786). ^13^ Tumor growth was monitored by measuring tumor size using calipers once or twice weekly, and tumor volume was calculated using the formula [length/2] × [width^2^]. When tumors reached a volume of 200-300mm^3^, 0.5 mg/mL of doxycycline (Sigma-Aldrich, D9891) was administered via drinking water for 2 weeks, with water refreshed every 3 to 4 days. To evaluate anti-tumor efficacy, tumor growth inhibition (%) was calculated using the formula (1 − [change of tumor volume in doxycycline-treated group ÷ change of tumor volume in doxycycline-untreated group]) × 100.

For combinatorial drug treatment studies, 2 × 10^6^ cells of ARID1A-mutant (Q456*/Q456*) cells or the surgically resected tissues from a patient with colon cancer with an ARID1A mutation (S764fs) were inoculated subcutaneously into the right flank of NSG mice. Tumor size was monitored as described earlier. When tumors reached approximately 100 mm^3^, mice were treated with 20 mg/kg of NSC617145 (TOCRIS, 5340), a selective WRN inhibitor, diluted in 20% (2-hydroxypropyl)-β-cyclodextrin (Sigma-Aldrich, C0926), administered intraperitoneally, and 30 mg/kg of UC2288 (Abcam, ab146969), a p21 inhibitor, diluted in a solution of 10% ethanol, 30% polyethylene glycol 400 (Sigma-Aldrich, 202398), and 60% PHOSAL 50 PG (Lipoid, 368315), administered orally. Both drugs were administered simultaneously every 2 days for a total of seven doses. Two days after the final treatment, tumor tissues were harvested and prepared for immunohistochemistry.

### Immunohistochemistry (IHC)

Tissue samples were fixed in 4% paraformaldehyde, dehydrated through a graded ethanol series, and embedded in paraffin. Sections of 4 μm in thickness were cut using a microtome and mounted on glass slides. Immunohistochemistry staining was performed using the BOND Polymer Refine Detection system (Leica, DS9800) in accordance with the manufacturer’s instructions. Following deparaffinization and rehydration, antigen retrieval was conducted using heat-induced epitope retrieval in ER1 (citrate buffer, pH 6.0), or in ER2 (EDTA buffer, pH 9.0). After washing, sections were incubated with primary antibodies at room temperature for 15 minutes. Sections were then incubated with Compact Polymer at room temperature for 8 minutes. The antigen-antibody complexes were visualized using 3,3′-diaminobenzidine. Sections were counterstained with hematoxylin, mounted with coverslips, and imaged at 400× magnification using an Aperio AT2 scanner (Leica). Antibody details and dilution ratios are provided in Supplementary Information 2.

### Statistical analysis

Statistical analyses were performed using GraphPad Prism (GraphPad Software, version 8.0.1) and R (version 4.4.2). Data are presented as mean ± standard deviation (SD) or mean ± standard error of the mean (SEM), as indicated in the figure legends. Differences between two groups were assessed using Welch’s unpaired t-test. For multiple-group comparisons, one-way or two-way analysis of variance (ANOVA) was performed, followed by Tukey’s, Dunnett’s, or Sidak’s post hoc tests, as specified in the figure legends.

## Acknowledgments

This work was supported by National Research Foundation of Korea grants (grant No. RS-2021-NR059767 and RS-2023-00222687) funded by the Ministry of Science and ICT (MSIT), Republic of Korea; by a grant from the Korean Health Technology R&D Project through the Korea Health Industry Development Institute (KHIDI), funded by the Ministry of Health and Welfare, Republic of Korea (grant No. RS-2024-00439165); and by a grant from the Creative-Pioneering Researchers Program through Seoul National University (grant No. 800-20200510). Schematics were created with bioRender (https://www.biorender.com).

## Author contributions

J.K. conceived and designed the study and performed most of the experiments. J.O., D.J., S.S., S.-J.L., S.E.L., Y.Y., D.K., A.-J.C., H.R.J., and Y.O. contributed to project discussions and provided feedback. S.-Y.C. supervised the study and wrote the manuscript. All authors reviewed and approved the final version of the manuscript.

