## Extended Data Figures for "Differential DNA damage response to WRN inhibition identifies a targetable vulnerability in ARID1A-mutated cancers"

#### Extended Data Fig. 1

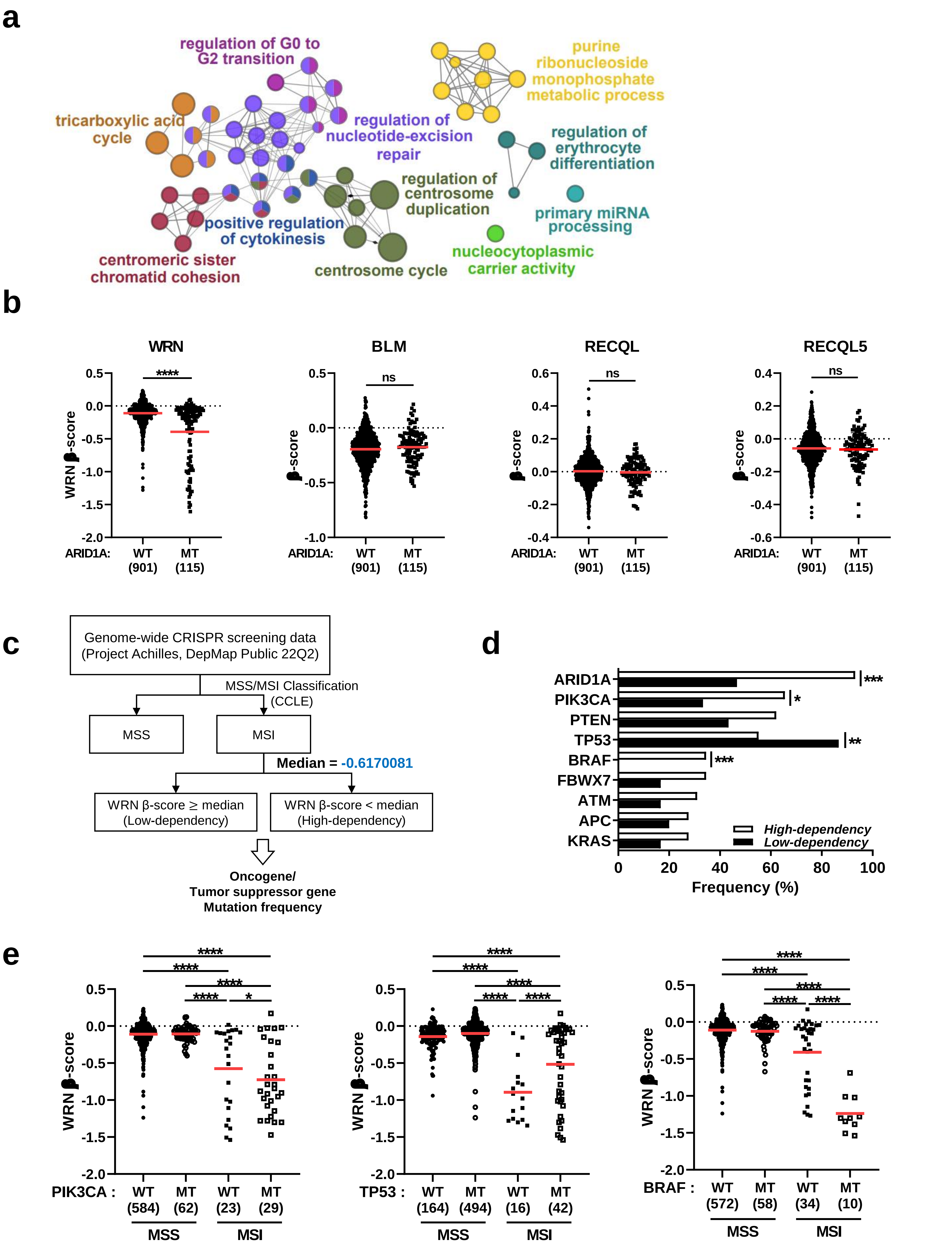

**Extended Data Fig. 1 Characterization of WRN dependency and associated genetic features in cancer cells**

(a) ClueGO enrichment analysis of genes with differential dependency scores based on Gene Ontology (GO) Biological Process gene sets. Nodes represent GO terms, with node size indicating enrichment significance. Functionally related terms are connected by edges and distinct functional categories are color-coded and labeled.

(b) Comparative analysis of β-scores for RecQ helicase family members (*WRN*, *BLM*, *RECQL*, and *RECQL5*) between ARID1A wild-type (n = 906) and ARID1A mutant (n = 115) cell lines. A red line indicates the mean β-score for each group. Statistical significance was determined using Welch's unpaired t-test.

(c) Schematic representation of the analysis using Project Achilles 22Q2 data, categorized by MSS and MSI status, followed by classification based on WRN dependency (High-dependency, β-scores < median; Low-dependency, β-scores ≥ median).

(d) Comparison of oncogene and tumor suppressor gene mutation frequencies between WRN high-dependency (n = 29) and WRN low-dependency (n = 30) groups. Statistical significance was determined using Fisher's exact test.

(e) Comparative analysis of β-scores for *WRN* in MSS and MSI cell lines, further classified by *PIK3CA* (left), *TP53* (middle), and *BRAF* (right) mutation status. Sample numbers are indicated on x-axis of each plot. A red line indicates the mean β-score for each group. Statistical significance was determined using Tukey's post hoc test following one-way analysis of variance (ANOVA).

Statistical significance was defined as follows: not significant; ns, $p > 0.05$; *, $p \leq 0.05$; **, $p \leq 0.01$; ***, $p \leq 0.001$; ****, $p \leq 0.0001$.

#### Extended Data Fig. 2

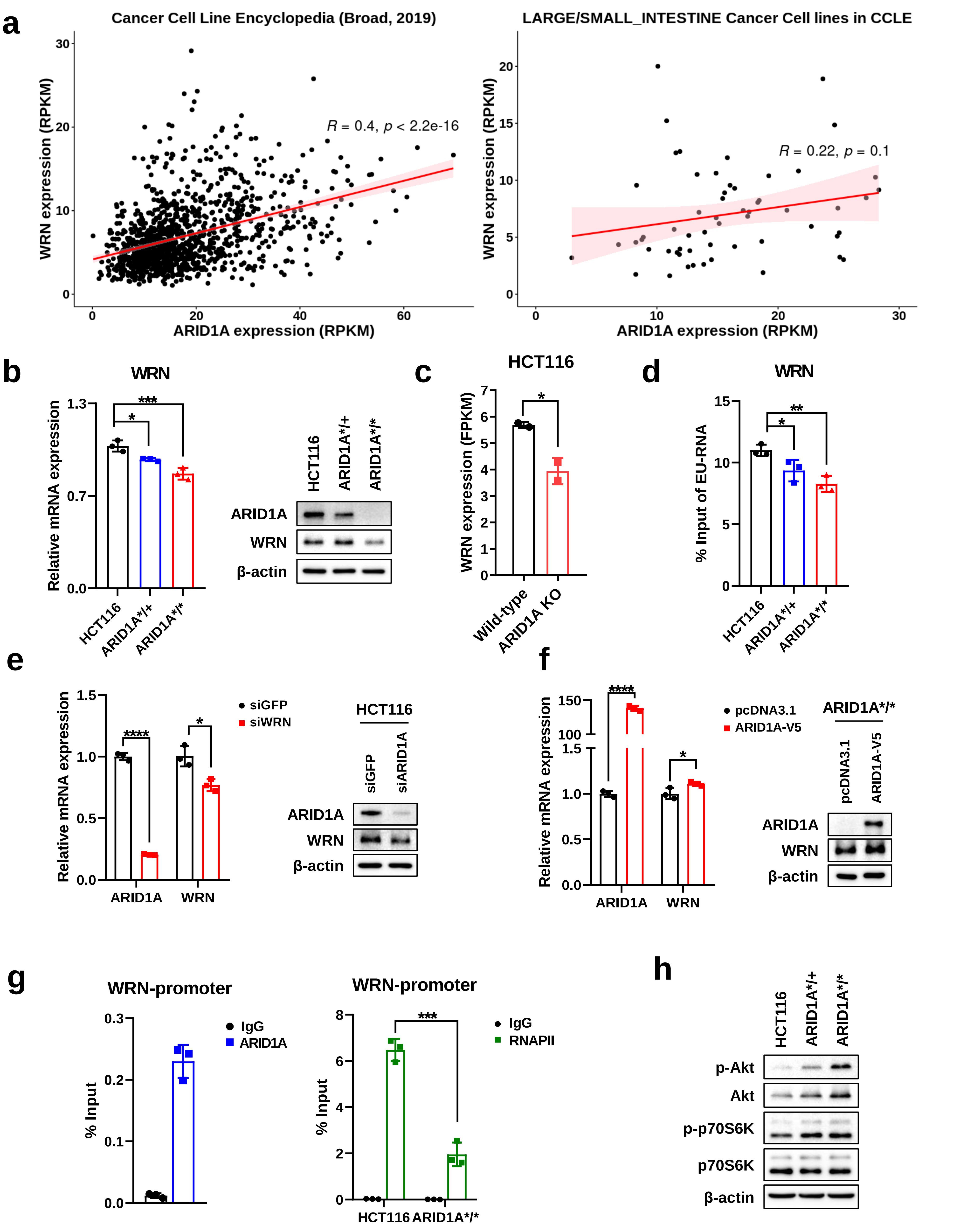

**Extended Data Fig. 2 ARID1A transcriptionally regulates WRN expression**

(a) Correlation plot of *ARID1A* and *WRN* mRNA expression levels in all cancer types (left, n = 1156) and large/small intestine cancer cell lines (right, n = 58) from the Cancer Cell Line Encyclopedia (CCLE). A red line represents the linear regression fit, and the light red shaded region indicates the confidence interval. Correlation analysis was assessed using Pearson correlation analysis.

(b) Quantitative RT-PCR data and western blot analysis of basal *WRN* expression levels in ARID1A wild-type and ARID1A-mutated cells. Data represent three independent experiments and are presented as mean ± standard deviation (SD). Statistical significance was determined using Welch's unpaired t-test.

(c) *WRN* mRNA expression levels of ARID1A wild-type and ARID1A-knockout (KO) HCT116 cells from GEO dataset GSE101966. Data represent two independent experiments and are presented as mean ± SD. Statistical significance was determined using Welch's unpaired t-test.

(d) Nascent RNA expression analysis in ARID1A wild-type and ARID1A-mutated cells. Nascent RNA was captured by incubating cells with 5-ethynyl uridine (EU, 0.5 mM) for 1 hour. Data represent three independent experiments and are presented as mean ± SD. Statistical significance was determined using Dunnett's post hoc test following one-way ANOVA.

(e,f) Quantitative RT-PCR and western blot analysis of *WRN* expression following ARID1A knockdown in ARID1A wild-type HCT116 cells (e) and ARID1A overexpression in ARID1A-mutated HCT116 cells (f). Data represent three independent experiments and are presented as mean ± SD. Statistical significance was determined using Welch's unpaired t-test.

(g) Chromatin immunoprecipitation (ChIP)-qPCR analysis of ARID1A and RNA polymerase II (RNAPII) binding at *WRN* promoter. Data represent three independent experiments and are presented as mean ± SD. Statistical significance was determined using Welch's unpaired t-test.

(h) Western blot analysis of AKT pathway in ARID1A wild-type and ARID1A-mutated cells.

Statistical significance was defined as follows: not significant; ns, $p > 0.05$; *, $p \leq 0.05$; **, $p \leq 0.01$; ***, $p \leq 0.001$; ****, $p \leq 0.0001$. RPKM, Reads Per Kilobase of transcript, per Million mapped reads.

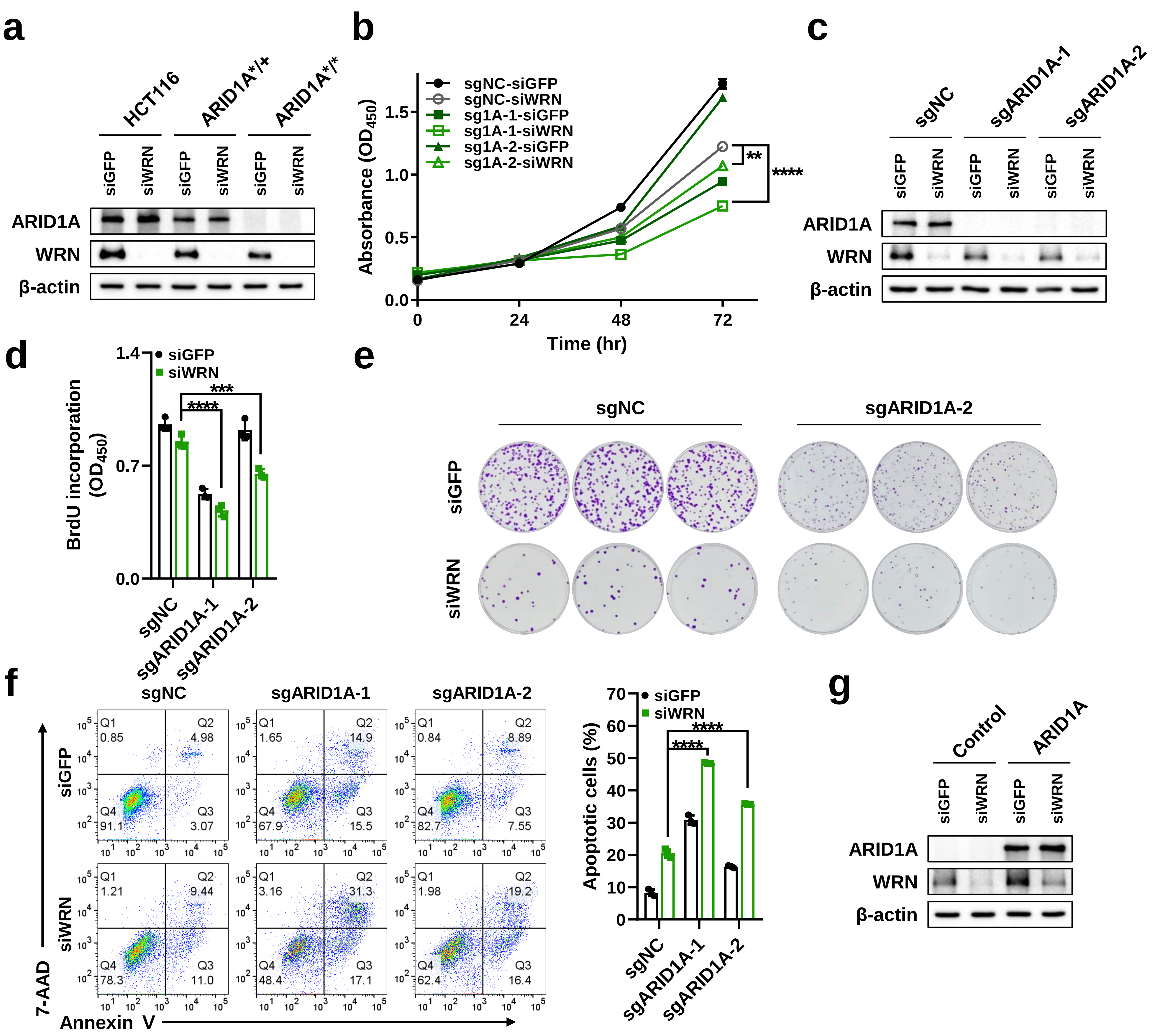


**Extended Data Fig. 3 Effect of WRN depletion on proliferation and apoptosis in ARID1A-knockout cells**

(a) Western blot analysis of *WRN* knockdown in ARID1A wild-type and ARID1A-mutated cells.

(b) Cell proliferation measured using the WST-1 assay in ARID1A wild-type and ARID1A- knockout HCT116 cells following *WRN* knockdown, measured at 0, 24, 48, and 72 hours after transfection. Data represent three independent experiments and are presented as mean ± SD. Statistical significance was determined using Tukey's post hoc test following one-way ANOVA. sgNC, Non-target control sgRNA.

(c) Western blot analysis of *WRN* knockdown in ARID1A wild-type and ARID1A-knockout HCT116 cells.

(d) BrdU incorporation assay to assess DNA synthesis in ARID1A wild-type and ARID1A-knockout HCT116 cells following WRN knockdown. Cells were incubated with BrdU for 8 hours at 24 hours after transfection. Data represent three independent experiments and are presented as mean ± SD. Statistical significance was determined using Tukey's post hoc test following one-way ANOVA.

(e) Representative images of the colony formation assay in ARID1A wild-type and ARID1A- knockout HCT116 cells following *WRN* knockdown, assessed after 9 days of culture.

(f) Apoptosis assay in ARID1A wild-type and ARID1A-knockout HCT116 cells following *WRN* knockdown, measured by annexin V/7-AAD staining at 72 hours after transfection. Representative flow cytometry plots (left) and quantification (right) are shown. Data represent three independent experiments and are presented as mean ± SD. Statistical significance was determined using Tukey's post hoc test following one-way ANOVA

(g) Western blot analysis of *WRN* knockdown in control and ARID1A-overexpressing ARID1A-mutated HCT116 cells.

Statistical significance was defined as follows: not significant; ns, $p > 0.05$; *, $p \leq 0.05$; **, $p \leq 0.01$; ***, $p \leq 0.001$; ****, $p \leq 0.0001$.

### Extended Data Fig. 4

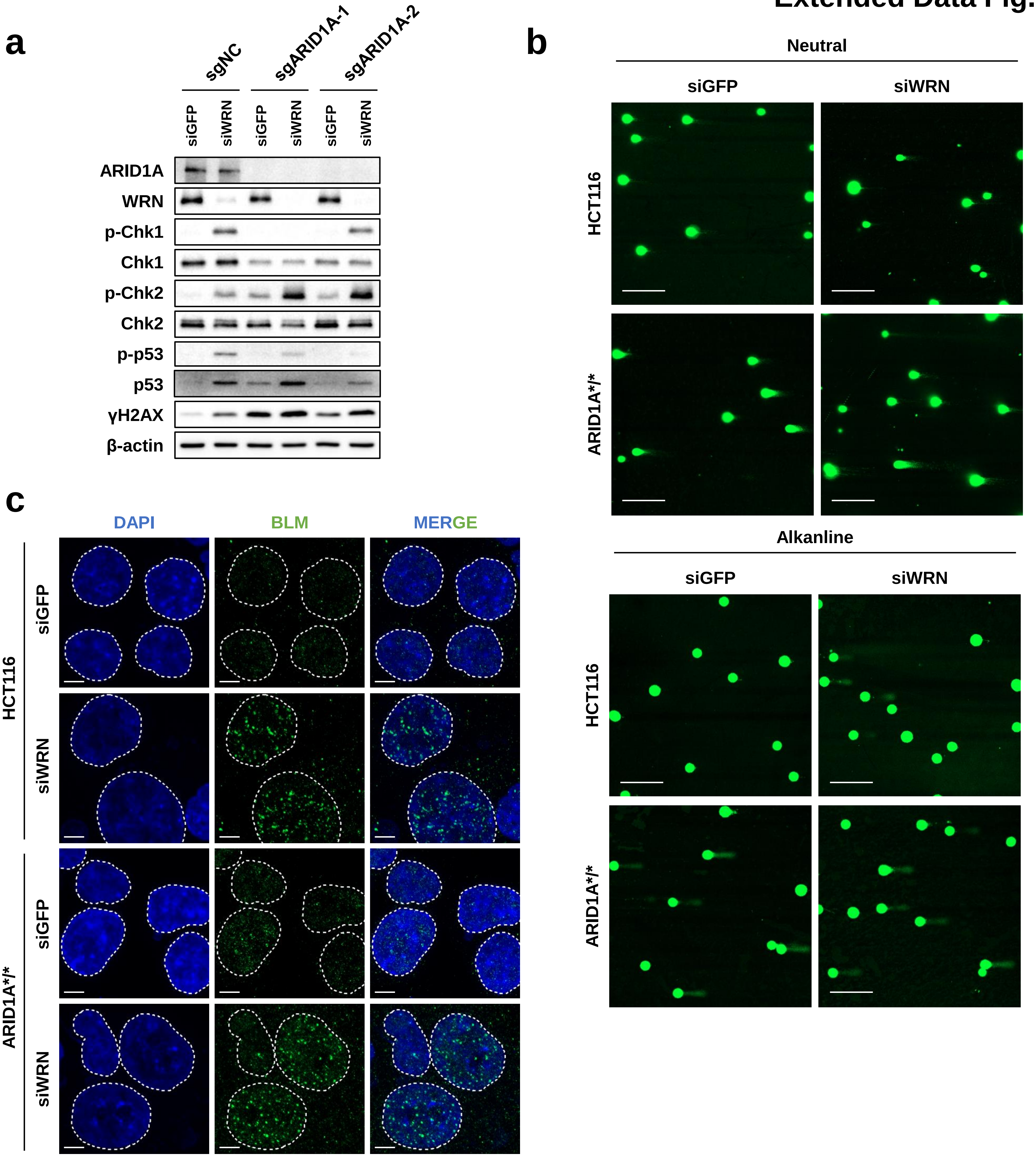


**Extended Data Fig. 4 DNA damage response upon WRN depletion in ARID1A-deficient cells**

(a) Western blot analysis of cell cycle regulatory proteins following *WRN* knockdown in ARID1A wild-type and ARID1A-knockout HCT116 cells.

(b) Comet assay images for neutral (top) and alkaline (bottom) comet assays following *WRN* knockdown in ARID1A wild-type and ARID1A-mutated HCT116 cells. Scale bar, 200 µm.

(c) Immunofluorescence images of BLM foci following *WRN* knockdown in ARID1A wild-type and ARID1A-mutated cells. White dotted lines indicate nuclear outlines. Scale bar, 5 µm.

### Extended Data Fig. 5

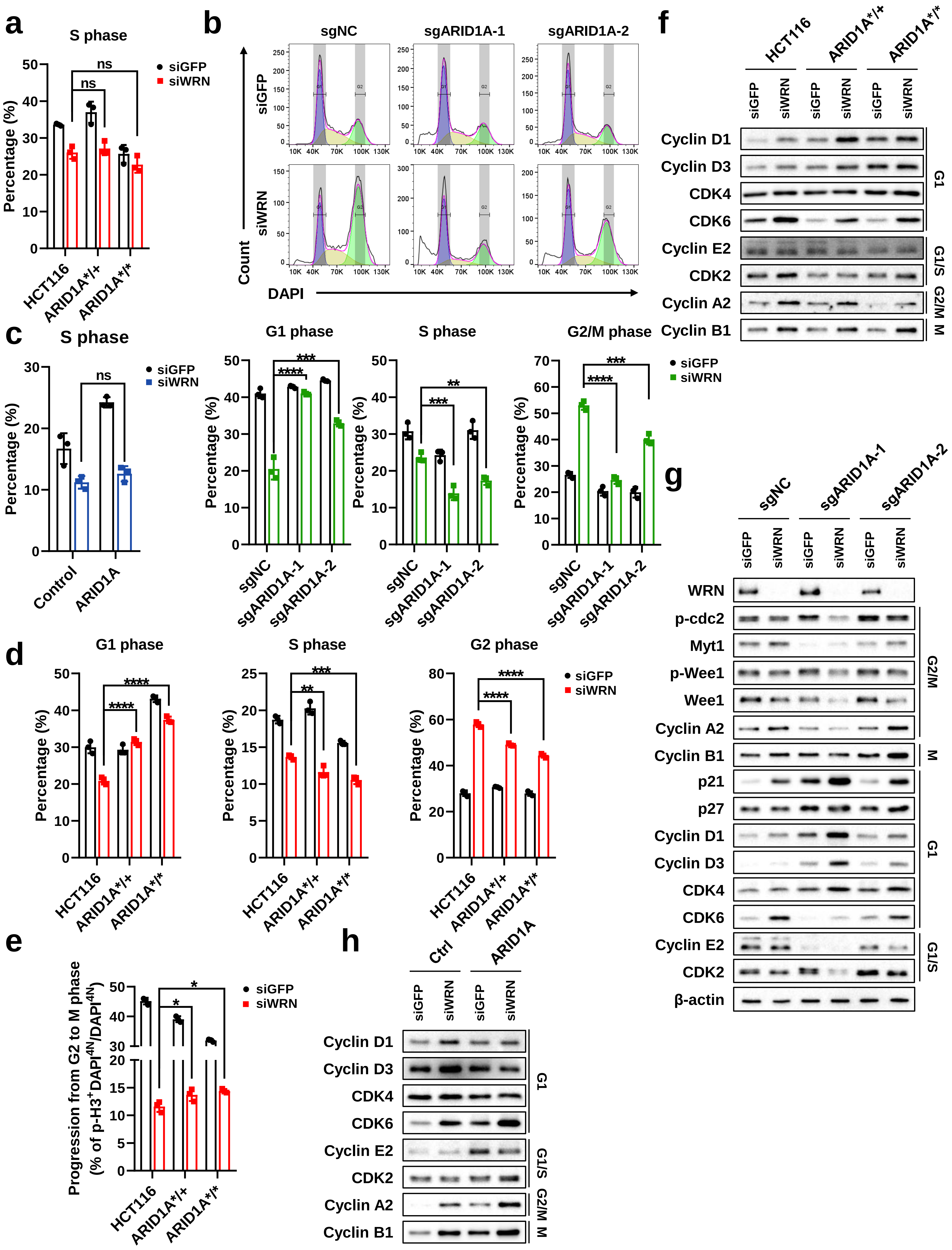

**Extended Data Fig. 5 Extended analysis of WRN depletion effects on cell cycle regulation**

(a) S phase proportion from cell cycle analysis following *WRN* knockdown in ARID1A wild-type and ARID1A-mutated cells. Data represent three independent experiments and are presented as mean ± SD. Statistical significance was determined using Tukey's post hoc test following one-way ANOVA.

(b) Cell cycle analysis following *WRN* knockdown in ARID1A wild-type and ARID1A-knockout HCT116 cells. Representative flow cytometry plots (top) and quantification (bottom) are shown. Data represent three independent experiments and are presented as mean ± SD. Statistical significance was determined using Tukey's post hoc test following one-way ANOVA.

(c) S phase proportion from cell cycle analysis following *WRN* knockdown in control and ARID1A-overexpressing ARID1A-mutated HCT116 cells. Data represent three independent experiments and are presented as mean ± SD. Statistical significance was determined using Tukey's post hoc test following one-way ANOVA.

(d) G1, S, and G2 phase distribution of phospho-histone H3 (p-H3) and DAPI staining following *WRN* knockdown in ARID1A wild-type and ARID1A-mutated cells. Cells were treated with paclitaxel (10 nM, 5 hours) at 48 hours after transfection. Data represent three independent experiments and are presented as mean ± SD. Statistical significance was determined using Tukey's post hoc test following one-way ANOVA.

(e) Proportion of cells progressing from G2 to M phase based on p-H3 staining assay following *WRN* knockdown in ARID1A wild-type and ARID1A-mutated cells. The proportion of p-H3-positive cells among total G2/M is considered as cells progressing from G2 to M phase. Data represent three independent experiments and are presented as mean ± SD. Statistical significance was determined using Tukey's post hoc test following one-way ANOVA.

(f-h) Western blot analysis of cell cycle regulatory proteins following *WRN* knockdown in ARID1A wild-type and ARID1A-mutated cells (f), in ARID1A wild-type and ARID1A-knockout HCT116 cells (g), and in control and ARID1A-overexpressing ARID1A-mutated HCT116 cells (h).

Statistical significance was defined as follows: not significant; ns, $p > 0.05$; *, $p \leq 0.05$; **, $p \leq 0.01$; ***, $p \leq 0.001$; ****, $p \leq 0.0001$.

### Extended Data Fig. 6

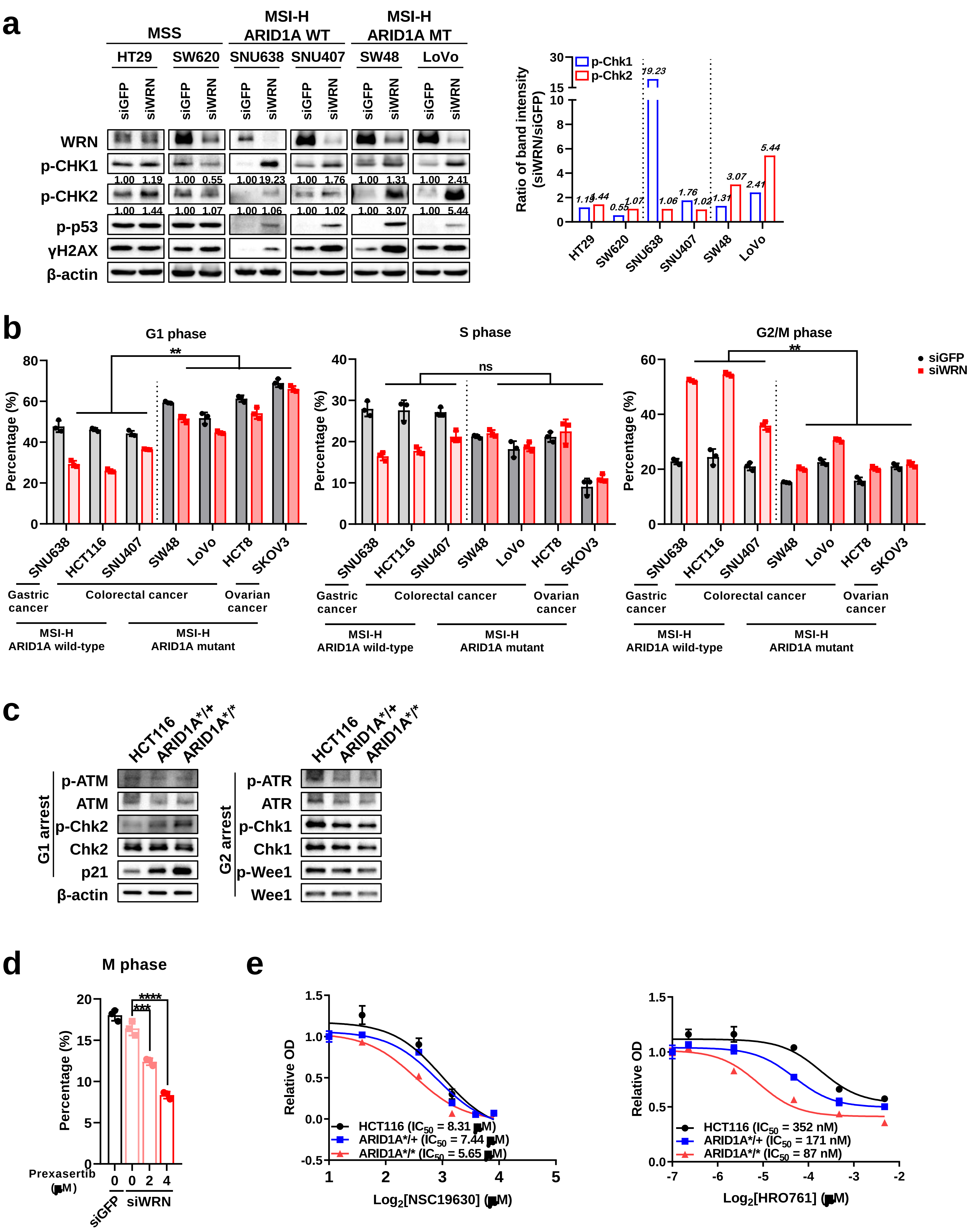

**Extended Data Fig. 6 DNA damage response and cell cycle analysis in MSI-H and ARID1A-mutant cell lines**

(a) Western blot analysis of DNA damage response proteins in MSS, MSI-H/ARID1A wild-type, and MSI-H/ARID1A mutant cell lines. The quantified siWRN/siGFP band intensity ratio for p-Chk1 and p-Chk2 are displayed below the blot and represented in the accompanying bar plot on the right.

(b) Cell cycle analysis of MSI-H/ARID1A wild-type and MSI-H/ARID1A mutant cell lines.

Data represent three independent experiments and are presented as mean ± SD. Statistical significance was determined using the Nested t-test.

(c) Western blot analysis of basal DNA damage response protein expression in HCT116 wild-type and ARID1A-mutated cells.

(d) M phase proportion from phospho-histone H3 staining assay in ARID1A wild-type HCT116 cells upon *WRN* knockdown (6 hours), followed by prexasertib treatment (24 hours). Cells were treated with paclitaxel (10 nM, 5 hours) at 24 hours after prexasertib treatment. Data represent three independent experiments and are presented as mean ± SD. Statistical significance was determined using Tukey's post hoc test following one-way ANOVA.

(e) Drug cytotoxicity assay with NSC19630 (left, 72 hours) and HRO761 (right, 72 hours) treatment in ARID1A wild-type and ARID1A-mutated cells. Drug concentrations ranged from 0 to 15 μM for NSC19630 and 0 to 5 μM for HRO761. $IC_{50}$ values are displayed on the plot with a fitted dose-response curve.

Statistical significance was defined as follows: not significant; ns, $p > 0.05$; *, $p \leq 0.05$; **, $p \leq 0.01$; ***, $p \leq 0.001$; ****, $p \leq 0.0001$.

### Extended Data Fig. 7

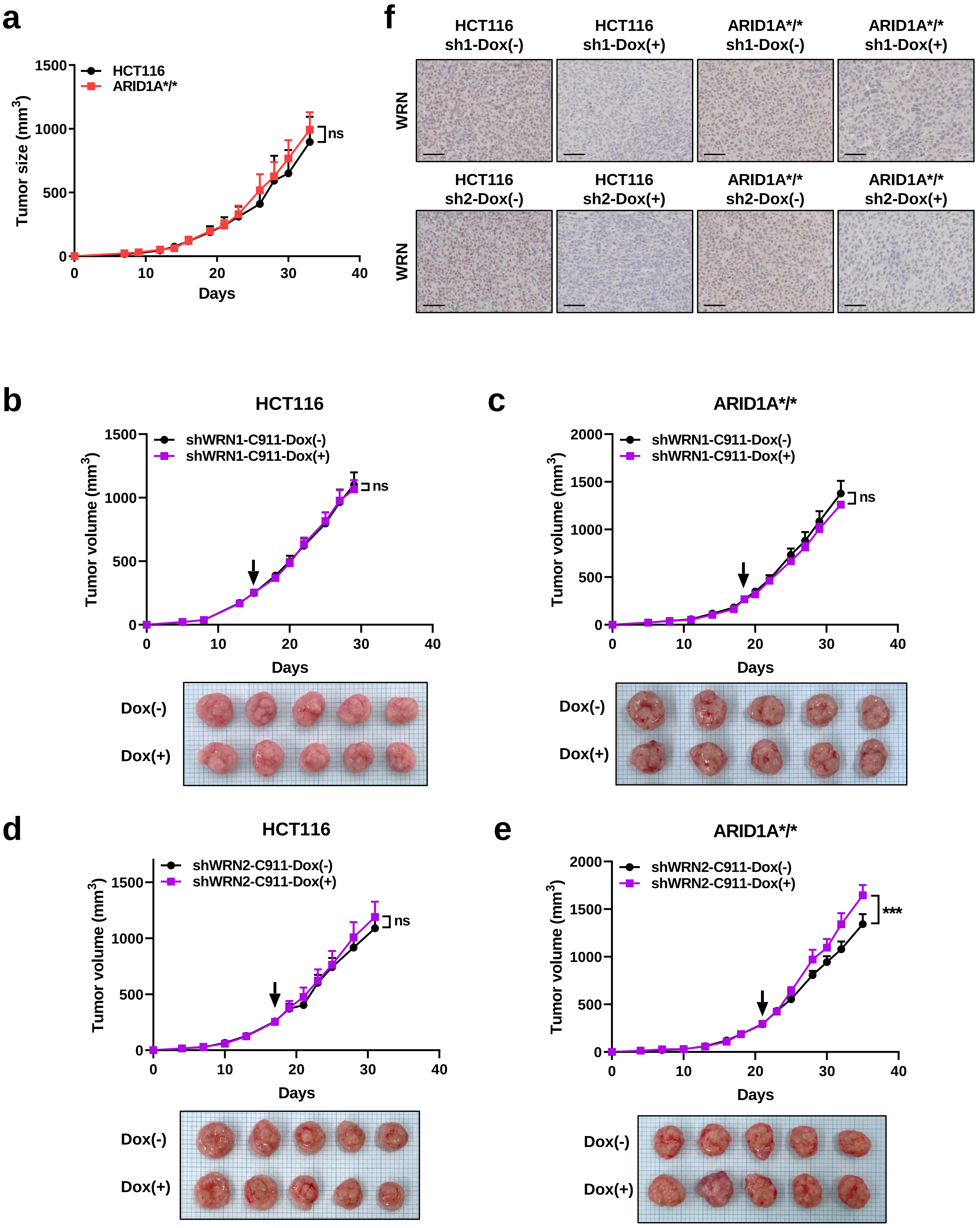

**Extended Data Fig. 7 Tumor growth analysis of WRN depletion controls using C911 shRNA in vivo**

(a) Tumor growth curves of xenografts using ARID1A wild-type and ARID1A-mutated HCT116 cells in NSG mice (n = 3 per group). Data are presented as mean ± standard error of the mean (SEM) and statistical significance was determined using Sidak's post hoc test following two-way ANOVA.

(b,c) Tumor growth curves for shWRN1-C911 xenografts with ARID1A wild-type HCT116 (b) and ARID1A-mutated HCT116 cells (c) in NSG mice (top, n = 5 per group). The arrow indicates the start of doxycycline (DOX) treatment. Representative tumor images from sacrificed mice at the end of the experiment are shown (bottom). Data are presented as mean ± SEM and statistical significance was determined using Sidak's post hoc test following two-way ANOVA.

(d,e) Tumor growth curves for shWRN2-C911 xenografts with ARID1A wild-type HCT116 (d) and ARID1A-mutated HCT116 cells (e) in NSG mice (top, n = 5 per group). The arrow indicates the start of DOX treatment. Representative tumor images from sacrificed mice at the end of the experiment are shown (bottom). Data are presented as mean ± SEM and statistical significance was determined using Sidak's post hoc test following two-way ANOVA.

(g) Representative immunohistochemistry images of WRN staining in tumors from shWRN1 and shWRN2 xenografts. Scale bar, 50 μm.

Statistical significance was defined as follows: not significant; ns, $p > 0.05$; *, $p \leq 0.05$; **, $p \leq 0.01$; ***, $p \leq 0.001$; ****, $p \leq 0.0001$.

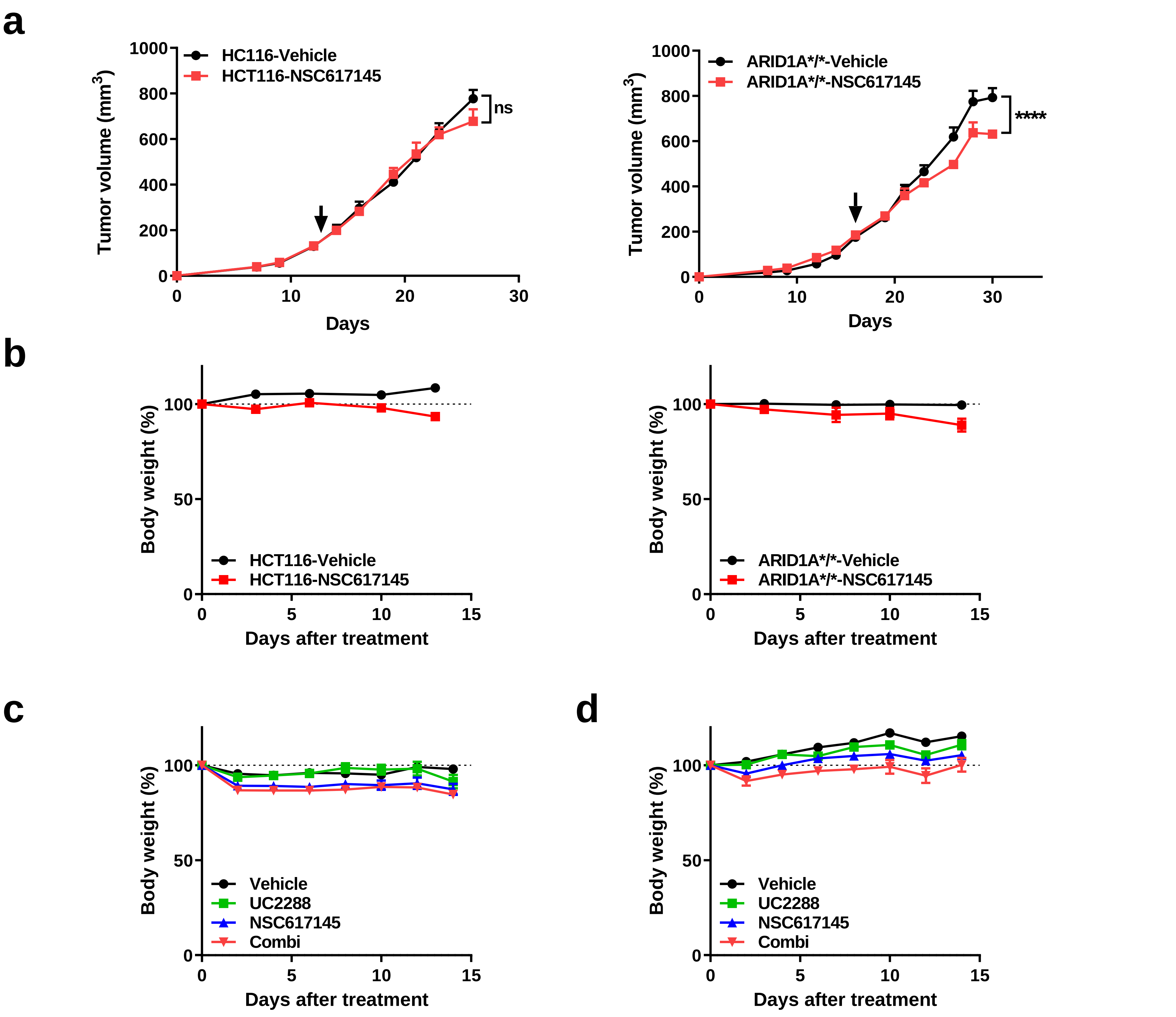


**Extended Data Fig. 8 Tumor growth analysis following NSC617145 treatment in vivo**

(a) Tumor growth curves for xenografts with ARID1A wild-type HCT116 cells (left, n = 5 for each group) and ARIDA-mutated HCT116 cells (right, n = 5 for vehicle, 4 for NSC617145) with NSC617145 treatment (20 mg/kg). The arrow indicates the start of NSC617145 treatment. Data are presented as mean ± standard error of the mean (SEM) and statistical significance was determined using Sidak's post hoc test following two-way analysis of variance (ANOVA).

(b) Body weight measurements over time following NSC617145 treatment. Data are presented as mean ± SEM.

(c,d) Body weight measurement over time following NSC617145 (20 mg/kg) and UC2288 (30 mg/kg) treatment in ARID1A-mutated HCT116 cells xenograft models (c) and patient-derived xenograft models (d). Data are presented as mean ± SEM.

Statistical significance was defined as follows: not significant; ns, $p > 0.05$; *, $p \leq 0.05$; **, $p \leq 0.01$; ***, $p \leq 0.001$; ****, $p \leq 0.0001$.

### Extended Data Fig. 9

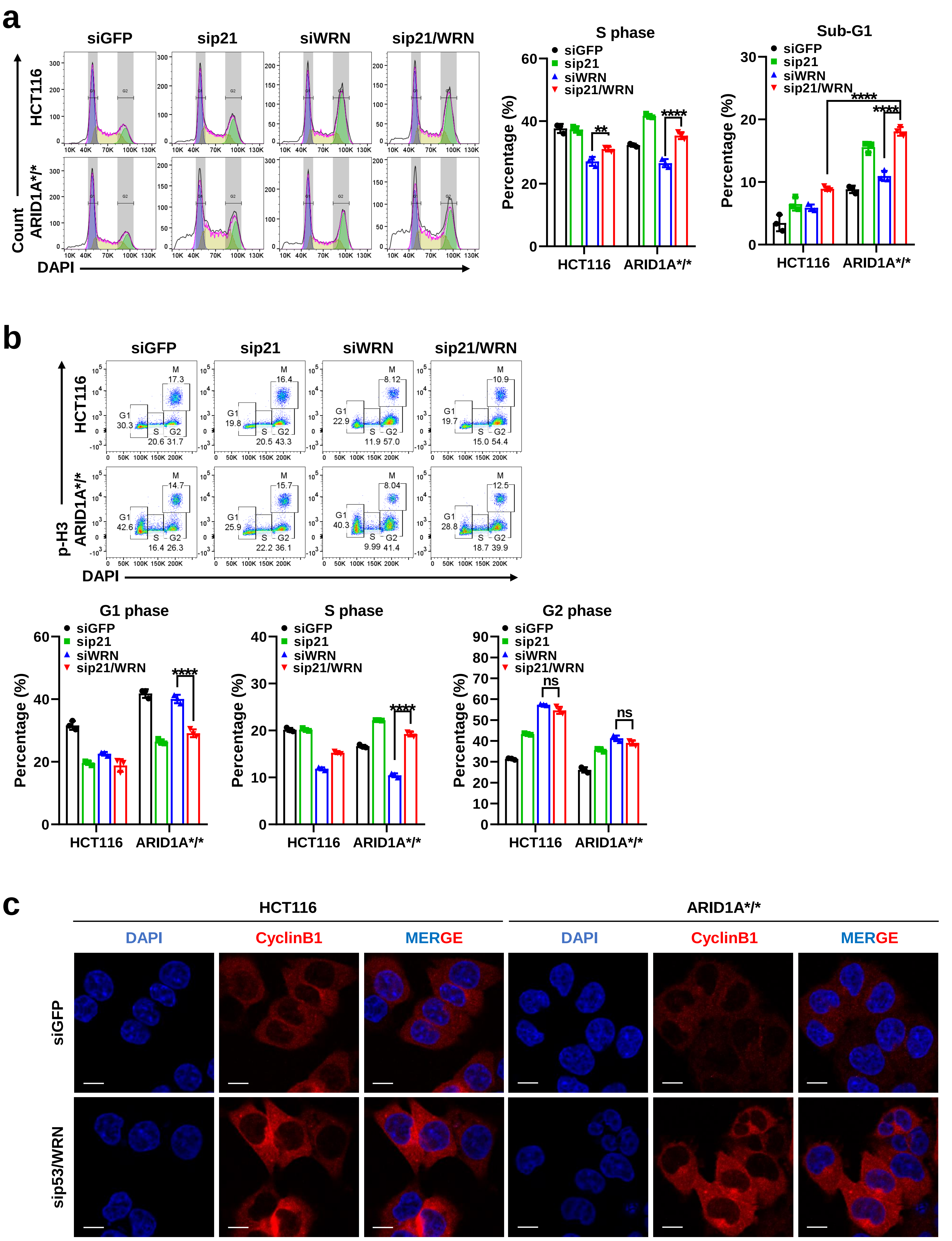

**Extended Data Fig. 9 Additional cell cycle distribution following combinatorial inhibition of WRN and p21**

(a) S phase and Sub-G1 phase proportions from cell cycle analysis following *WRN* and *p21* knockdown in ARID1A wild-type and ARID1A-mutated cells. Representative flow cytometry plots (left) and quantification (right) are shown. Data represent three independent experiments and are presented as mean ± SD. Statistical significance was determined using Dunnett's post hoc test following one-way ANOVA.

(b) G1, S, and G2 phase proportions from cell cycle analysis following *WRN* and *p21* knockdown in ARID1A wild-type and ARID1A-mutated cells. Representative flow cytometry plots (top) and quantification (bottom) are shown. Data represent three independent experiments and are presented as mean ± SD. Statistical significance was determined using Dunnett's post hoc test following one-way ANOVA.

(c) Immunofluorescence images of cyclinB1 and DAPI staining following *WRN* and *p53* knockdown in ARID1A wild-type and ARID1A-mutated cells. Scale bar, 10 μm.

Statistical significance was defined as follows: not significant; ns, $p > 0.05$; *, $p \leq 0.05$; **, $p \leq 0.01$; ***, $p \leq 0.001$; ****, $p \leq 0.0001$.

### Extended Data Fig. 10

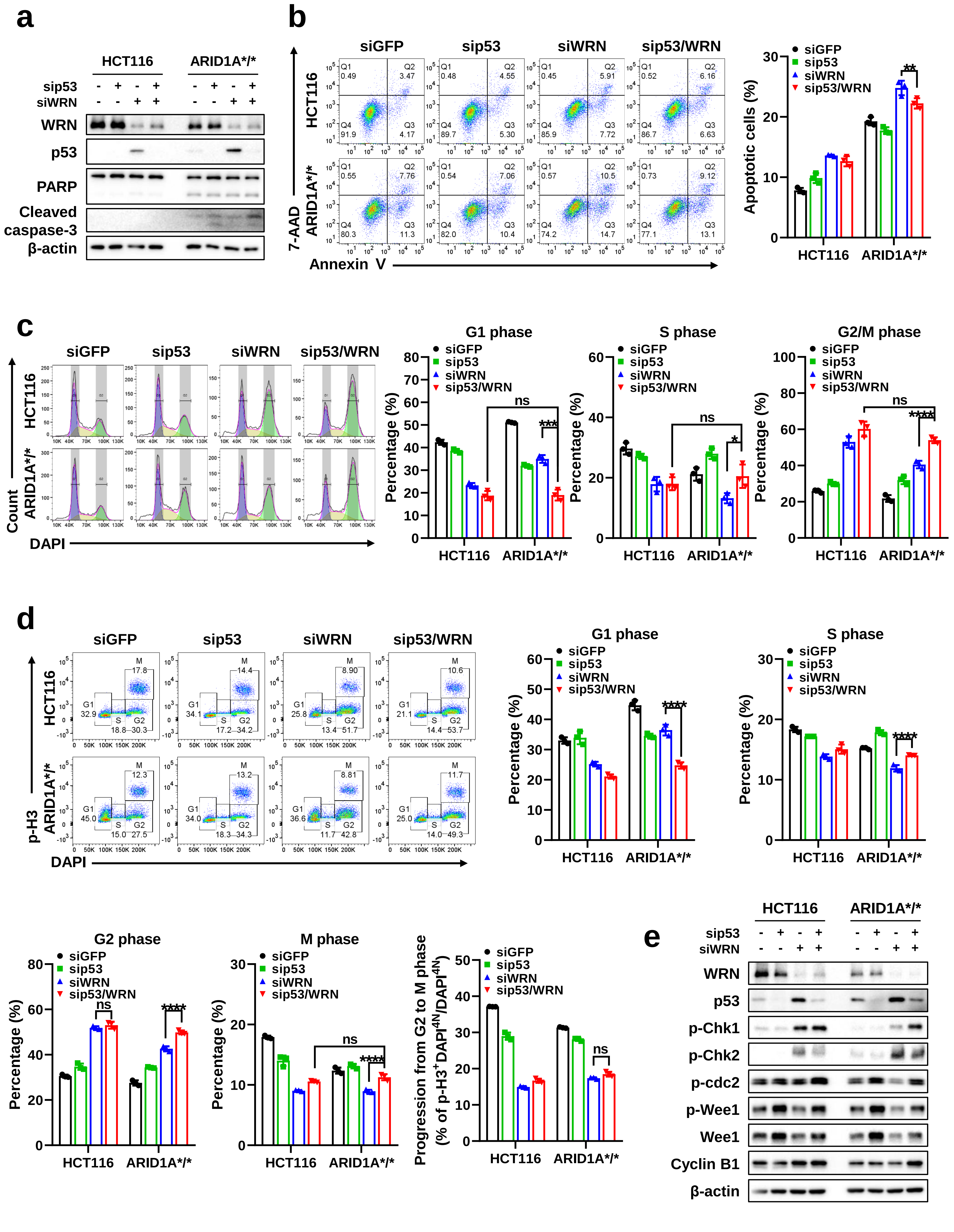

**Extended Data Fig. 10 Effect of WRN and p53 dual inhibition on cell cycle progression**

(a) Western blot analysis of apoptosis markers following *WRN* and *p53* knockdown in ARID1A wild-type and ARID1A-mutated cells.

(b) Apoptosis assay following *WRN* and *p53* knockdown in ARID1A wild-type and ARID1A-mutated cells, measured by annexin V/7-AAD staining at 48 hours after transfection. Representative flow cytometry plots (left) and quantification (right) are shown. Data represent three independent experiments and are presented as mean ± SD. Statistical significance was determined using Dunnett's post hoc test following one-way ANOVA.

(c) Cell cycle analysis following *WRN* and *p53* knockdown in ARID1A wild-type and ARID1A-mutated cells. Representative flow cytometry plots (left) and quantification (right) are shown. Data represent three independent experiments and are presented as mean ± SD. Statistical significance was determined using Dunnett's post hoc test following one-way ANOVA.

(d) Phospho-histone H3 staining assay following *WRN* and *p53* knockdown in ARID1A wild-type and ARID1A-mutated cells. Cells were treated with paclitaxel (10 nM, 5 hours) at 48 hours after transfection. The proportion of p-H3-positive cells among total G2/M is considered as cells progressing from G2 to M phase. Representative flow cytometry plots (left) and quantification (right and bottom) are shown. Data represent three independent experiment and are presented as mean ± SD. Statistical significance was determined using Dunnett's post hoc test following one-way ANOVA.

(e) Western blot analysis of cell cycle regulatory proteins following *WRN* and *p53* knockdown in ARID1A wild-type and ARID1A-mutated cells.

Statistical significance was defined as follows: not significant; ns, $p > 0.05$; *, $p \leq 0.05$; **, $p \leq 0.01$; ***, $p \leq 0.001$; ****, $p \leq 0.0001$.

**Supplementary Information 1. Information for qPCR primers**

| Gene | Forward | Reverse |
|---|---|---|
| **GAPDH** | CCAAAATCAAGTGGGGCGAT | TGCTGATGATCTTGAGGCTG |
| **ARID1A** | CCAGTAAGGGAGGGCAAGAA | CTCCTTGGCTGCTGGAAATC |
| **WRN** | TGCTAGTGATTGCTCTTTCCTG | CCTTGCCAAGTTTCCCTCTATTG |

**Supplementary Information 2. Information for antibodies**

| Antibodies for immunoblotting | | | |
|---|---|---|---|
| **Antibody Name** | **Company** | **Cat. No.** | **Dilution** |
| ARID1A | Cell Signaling Technology, Inc. | #12354 | 1:1000 |
| WRN | Cell Signaling Technology, Inc. | #4666 | 1:1000 |
| Phospho-Akt | Cell Signaling Technology, Inc. | #9271 | 1:1000 |
| Akt | Cell Signaling Technology, Inc. | #9272 | 1:1000 |
| Phospho-p70 S6 Kinase | Cell Signaling Technology, Inc. | #9205 | 1:1000 |
| p70 S6K Kinase | Cell Signaling Technology, Inc. | #9202 | 1:1000 |
| Phospho-ATM | Cell Signaling Technology, Inc. | #5883 | 1:500 |
| Phospho-ATR | Cell Signaling Technology, Inc. | #2853 | 1:1000 |
| Phospho-Chk1 | Cell Signaling Technology, Inc. | #2348 | 1:1000 |
| Chk1 | Santa Cruz Biotechnology, Inc. | sc-8408 | 1:1000 |
| Phospho-Chk2 | Cell Signaling Technology, Inc. | #2197 | 1:1000 |
| Chk2 | Cell Signaling Technology, Inc. | #6334 | 1:1000 |
| Phospho-p53 | Cell Signaling Technology, Inc. | #9286 | 1:500 |
| p53 | Cell Signaling Technology, Inc. | #2524 | 1:1000 |
| Phospho-Histone H2A.X | Cell Signaling Technology, Inc. | #9718 | 1:1000 |
| Cyclin D1 | Cell Signaling Technology, Inc. | #2978 | 1:1000 |
| Cyclin D3 | Cell Signaling Technology, Inc. | #2936 | 1:1000 |
| Cyclin E2 | Cell Signaling Technology, Inc. | #4132 | 1:1000 |
| Cyclin A2 | Cell Signaling Technology, Inc. | #4656 | 1:1000 |
| Cyclin B1 | Cell Signaling Technology, Inc. | #12231 | 1:1000 |
| CDK4 | Cell Signaling Technology, Inc. | #12790 | 1:1000 |
| CDK6 | Cell Signaling Technology, Inc. | #3136 | 1:1000 |
| CDK2 | Cell Signaling Technology, Inc. | #2546 | 1:1000 |
| Phospho-cdc2 | Cell Signaling Technology, Inc. | #4539 | 1:1000 |
| Myt1 | Cell Signaling Technology, Inc. | #4282 | 1:1000 |
| Phospho-Wee1 | Cell Signaling Technology, Inc. | #4910 | 1:1000 |
| Wee1 | Santa Cruz Biotechnology, Inc. | sc-5285 | 1:1000 |
| p18 INK4C | Cell Signaling Technology, Inc. | #2896 | 1:1000 |
| p21 Waf1/Cip1 | Cell Signaling Technology, Inc. | #2947 | 1:1000 |
| p27 Kip1 | Cell Signaling Technology, Inc. | #3686 | 1:1000 |
| PARP | Cell Signaling Technology, Inc. | #9542 | 1:1000 |
| Cleaved Caspase-3 | Cell Signaling Technology, Inc. | #9661 | 1:1000 |
| β-Actin | Sigma-Aldrich | A2228 | 1:5000 |

| Antibodies for immunofluorescence | | | |
|---|---|---|---|
| **Antibody Name** | **Company** | **Cat. No.** | **Dilution** |
| ARID1A | Abcam PLC. | ab182560 | 1:100 |
| BLM | Santa Cruz Biotechnology, Inc. | sc-365753 | 1:100 |
| FANCD2 | Santa Cruz Biotechnology, Inc. | sc-20022 | 1:50 |
| DAPI | Sigma-Aldrich | DUO82040 | |

| Antibodies for immunohistochemistry | | | |
|---|---|---|---|
| **Antibody Name** | **Company** | **Cat. No.** | **Dilution** |
| Ki-67 | Novus Biologicals | NBP2-19012 | 1:500 |
| WRN | Bethyl Laboratories, Inc. | IHC-00407 | 1:300 |
| p21 | Cell Signaling Technology, Inc. | #2947 | 1:400 |
| γH2AX | Abcam | ab11174 | 1:8000 |
| cleaved caspase-3 | Cell Signaling Technology, Inc. | #9661 | 1:2000 |
